# Intercellular interaction between FAP fibroblasts and CD150 inflammatory monocytes mediates fibro-stenosis in Crohn’s disease

**DOI:** 10.1101/2023.05.04.539383

**Authors:** Bo-Jun Ke, Saeed Abdurahiman, Francesca Biscu, Gabriele Dragoni, Sneha Santhosh, Veronica De Simone, Anissa Zouzaf, Lies van Baarle, Michelle Stakenborg, Veronika Bosáková, Yentl Van Rymenant, Emile Verhulst, Sare Verstockt, Gabriele Bislenghi, Albert Wolthuis, Jan Frič, Christine Breynaert, Andre D’Hoore, Pieter Van Der Veken, Ingrid De Meester, Bram Verstockt, Gert de Hertogh, Séverine Vermeire, Gianluca Matteoli

## Abstract

Crohn’s disease (CD) is marked by recurring intestinal inflammation and tissue injury, often resulting in fibro-stenosis and bowel obstruction, necessitating surgical intervention with high recurrence rates. To elucidate complex intercellular interactions leading to fibro-stenosis in CD, we analysed the transcriptome of cells isolated from the transmural ileum of CD patients, including a trio of lesions from each patient: non-affected, inflamed, and stenotic ileum samples, and compared them with samples from non-CD patients. Our computational analysis revealed that pro-fibrotic signals from a subset of monocyte-derived cells expressing CD150 induce a disease-specific fibroblast population, resulting in chronic inflammation and tissue fibrosis. The transcription factor TWIST1 was identified as a key modulator of fibroblast activation and extracellular matrix (ECM) production. Therapeutic inhibition of TWIST1 inhibits fibroblast activation, reducing ECM production and deposition. These findings suggest that the myeloid-stromal axis may offer a promising therapeutic target to prevent fibro-stenosis in CD.

## INTRODUCTION

Crohn’s disease (CD) is a chronic inflammatory disorder of the gastrointestinal tract. Although patients suffering from CD share common clinical characteristics, the natural disease course is rather heterogeneous, in which the disease behaviour can remain indolent or progress rapidly toward severe comorbidities.^1–3^ About 75% of CD patients eventually require surgery due to uncontrollable inflammation, and/or complications with penetrating or stricturing disease. Stricture formation in the ileum is the most important indication for surgery in up to 70-80% of CD patients within their lifetime.^4, 5^ Ileal stricturing is caused by excessive extracellular matrix (ECM) deposition and muscular hyperplasia upon successive cycles of inflammation and tissue repair, which progressively lead to fibrosis and bowel obstruction, hence the term fibro-stenosis.^6, 7^

Current CD therapies aim to suppress inflammation through the administration of corticosteroids, immunosuppressive agents, and/or biologicals such as anti-tumour necrosis factor (TNF) antibodies, anti-integrins, anti-IL-12/23 agents or JAK inhibitors.^8^ While these therapies lead to symptomatic disease remission in approximately 40% of patients, recurrent flares still result in cumulative tissue damage and remodelling of the gut wall.^9^ Indeed, as no anti-fibrotic drugs are currently available for CD patients, the incidence of fibro-stenosis, an irreversible condition that may necessitate surgical resection, remains high.^10^ Most of the existing research has relied on biopsies procured through ileo-colonoscopy to examine mucosal inflammation and cellular heterogeneity in inflammatory bowel disease.^11–16^ Consequently, these studies did not address transmural inflammation and were unable to comprehensively elucidate the aetiology of bowel remodelling and obstruction within the ileum and colon of patients afflicted with CD. In the present study, we performed single-cell RNA sequencing (scRNA-seq) to characterise full-thickness of transmural terminal ileum from fibro-stenotic CD patients undergoing ileocaecal resection. To fully characterise the different stages of lesions in CD during disease progression, we profiled a trio of samples from each patient: proximal non-affected ileum, inflamed ileum with ulceration, and stenotic ileum. Using this approach, we aimed to characterise cells in the deeper layers of the gut and dissect the molecular and cellular processes of intestinal fibrosis and the complex interactions between immune and stromal cells.

Our study extends the knowledge on cell heterogeneity in the transmural ileum of fibro-stenotic CD patients and highlights a novel intercellular interaction between immune cells and fibroblasts, driving fibrosis in CD. We found that pathogenic CD150 Inflammatory monocytes promote tissue remodelling and fibrosis by inducing the differentiation of FAP fibroblasts via the transcriptional regulator TWIST1, specifically during inflammation and stenosis. Overall, our findings suggest that targeting the myeloid-mesenchymal axis during inflammation could be an effective strategy for developing new therapies to prevent fibro-stenosis in patients with CD.

## RESULTS

### Uncover the heterogeneity of fibroblasts in fibro-stenotic CD

Despite CD being a transmural disease, the use of endoscopic sampling has been a persistent limitation in previous studies.^14–16^ Therefore, aimed to investigate the cellular landscape and intercellular interactions in the transmural ileum of CD patients to identify potential therapeutic targets. First, we classified transmural biopsies from resected ileal tissue within each patient based on macroscopic features. Then, we determined inflammatory and fibrotic activity in the proximal healthy margin, inflamed, and stenotic ileum of CD patients, and control non-CD healthy ileum from colorectal cancer (CRC) patients using Hematoxylin and eosin (H&E) stain and Masson’s trichrome staining (Figure S1A). To evaluate our sample classification microscopically, we modified a histopathologic scoring system based on previous studies to assess inflammation and fibro-stenosis in CD ileum (Supplemental item Table S1 and Table S2).^17, 18^ We observed moderate to severe degrees of inflammation and fibrosis, including fissuring ulceration and abscess in inflamed ileum and stenotic ileum (Figure S1B, left and middle). Although similar fibrosis features were detected in both inflamed and stenotic ileum, stenotic ileum had a significantly higher level of overall collagen deposition (Figure S1A, shown in blue) compared to inflamed ileum, 35.9% versus 28.9% respectively (*p*<0.05) (Figure S1B, right). Altogether, these findings confirmed that our method of classification was reliable to study the progression of tissue through inflammation to fibro-stenosis.

To uncover the complex interplay between immune and stromal cells during fibrosis in CD, we profiled the transcriptome of 169,205 cells from transmural terminal ileum of CD patients (n = 10, a trio of lesions from each; proximal as unaffected, inflamed, and stenotic ileum) and CRC patients (n = 5; control ileum) using scRNA-seq (Figure 1A, Supplemental item Table S3). A similar number of cells were profiled from each of the three CD lesions; Proximal = 33.78%, Inflamed = 32.48%, Stenotic = 33.77% (Figure S1C to S1E). Unsupervised clustering followed by annotation of the integrated gene expression data identified several clusters which were classified into 8 compartments, including T cells and ILCs (*CD3D, CD3E, CD4, CD8A, NKG7, GNLY, KLRB1*), B cells (*CD19, CD79A, MS4A1*), plasma cells (*IGKC, MZB1, JCHAIN*), myeloid cells (*CD68, LYZ*), epithelial cells (*EPCAM*), endothelial cells (*VWF, PECAM1*), enteric glial cells (*PLP1, S100B*) and mesenchymal cells (*LUM, PDGFRA, DCN*) (Figure 1B to 1D). To identify the major ECM-producing cell compartments, we compared expression of ECM core genes between each cell compartment. This comparison confirmed the mesenchymal compartment as the major source of ECM proteins during inflammation and stenosis (Figure S1F). Thus, we proceeded with a deeper characterisation of the mesenchymal compartment to address their heterogeneity in different disease states. Re-clustering of the mesenchymal compartment identified two clusters of mural cells: pericytes (*RGS5*) and contractile pericytes (*ADIRF*), one cluster of smooth muscle cells (*MYOCD, MYH11, ACTG2*) and seven clusters of fibroblasts: myofibroblasts (*SOX6, ACTA2*), ADAMDEC1 fibroblasts (*ADAMDEC1, CCL11*), ABL2 fibroblasts (*ABL2, PLIN2, CLDN1*), GREM1^-^CD34^+^ fibroblasts (*MFAP5, CD55*), GREM1^+^CD34^+^ fibroblasts (*GREM1, C3, C7*), FAP fibroblasts (*FAP, CD82, TWIST1, POSTN*) and a cluster of proliferating FAP fibroblasts (*MKI67, TOP2A, CENPF*) (Figure 2A and 2B). FAP fibroblasts and ABL2 fibroblasts were found to be unique to inflamed and stenotic ileum compared to control and proximal ileum (Figure 2C, and S2A). To identify the stromal cluster responsible for pathological ECM deposition, we developed a collagen module score utilising the core matrisome collagen gene signature from MatrisomeDB. This approach identified FAP fibroblasts as the primary ECM-producing cells among all mesenchymal subsets (Figure 2D). This was confirmed by Gene ontology (GO) enrichment analysis, which showed that FAP fibroblasts were significantly enriched in GO processes associated with extracellular matrix organisation (Figure 2E).

**Figure 1.**
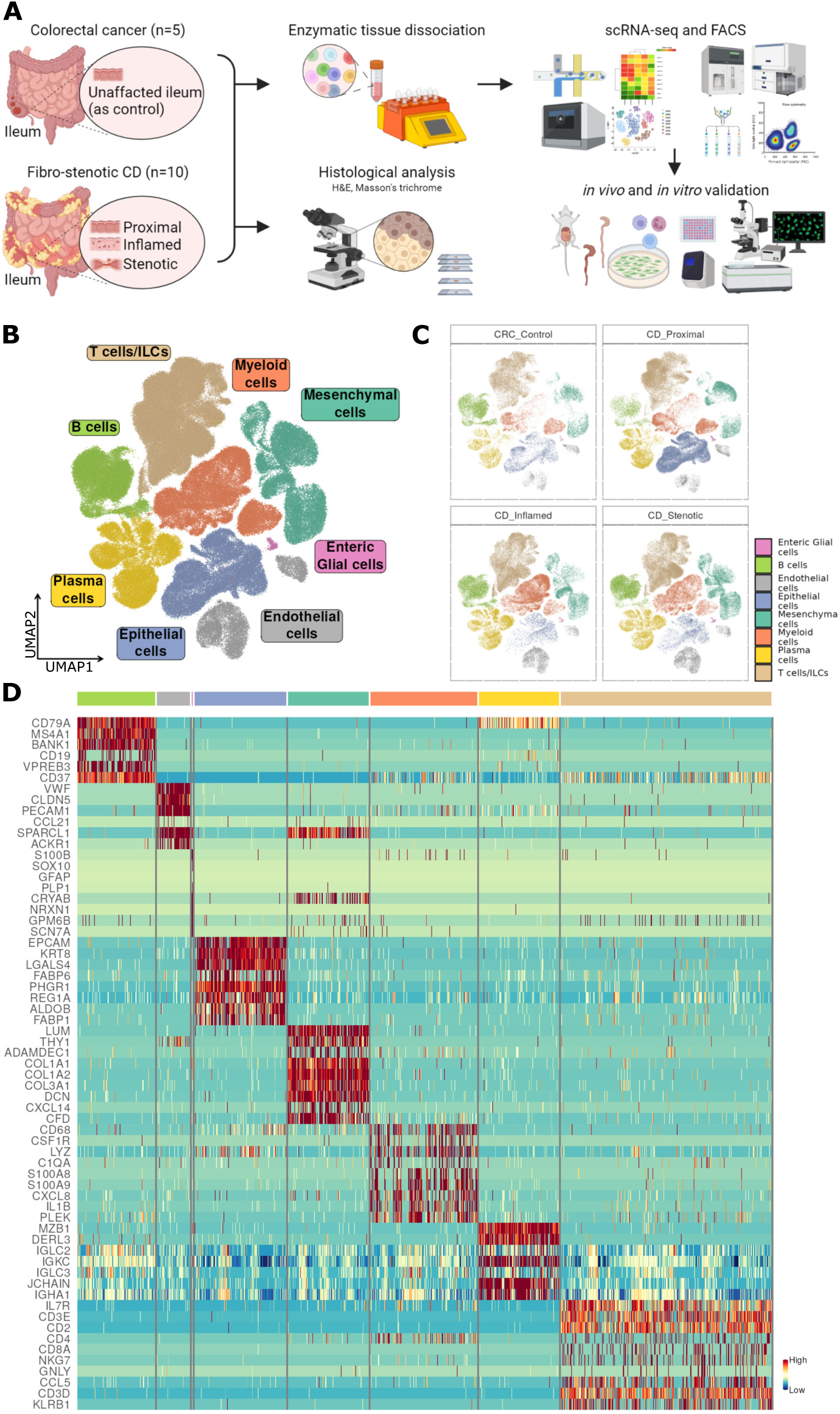
Single-cell profiling of fibro-stenotic ileal from CD and control ileum from CRC. (**A**) Experimental workflow for scRNA-seq of ileum using the 10x Chromium platform and further analyses and validations in this study. (**B)** Uniform Manifold Approximation and Projection (UMAP) embedding showing ileal single-cell transcriptomes from 169,547 cells from 10 CD patients with a trio of lesions (proximal, inflamed and stenotic) and 5 CRC control ileum depicting cell compartments. (**C**) UMAP in Figure 1B split by disease segments. (**D**) Heatmap depicting relative expression of distinguishing marker genes in each cell compartment.

**Figure 2.**
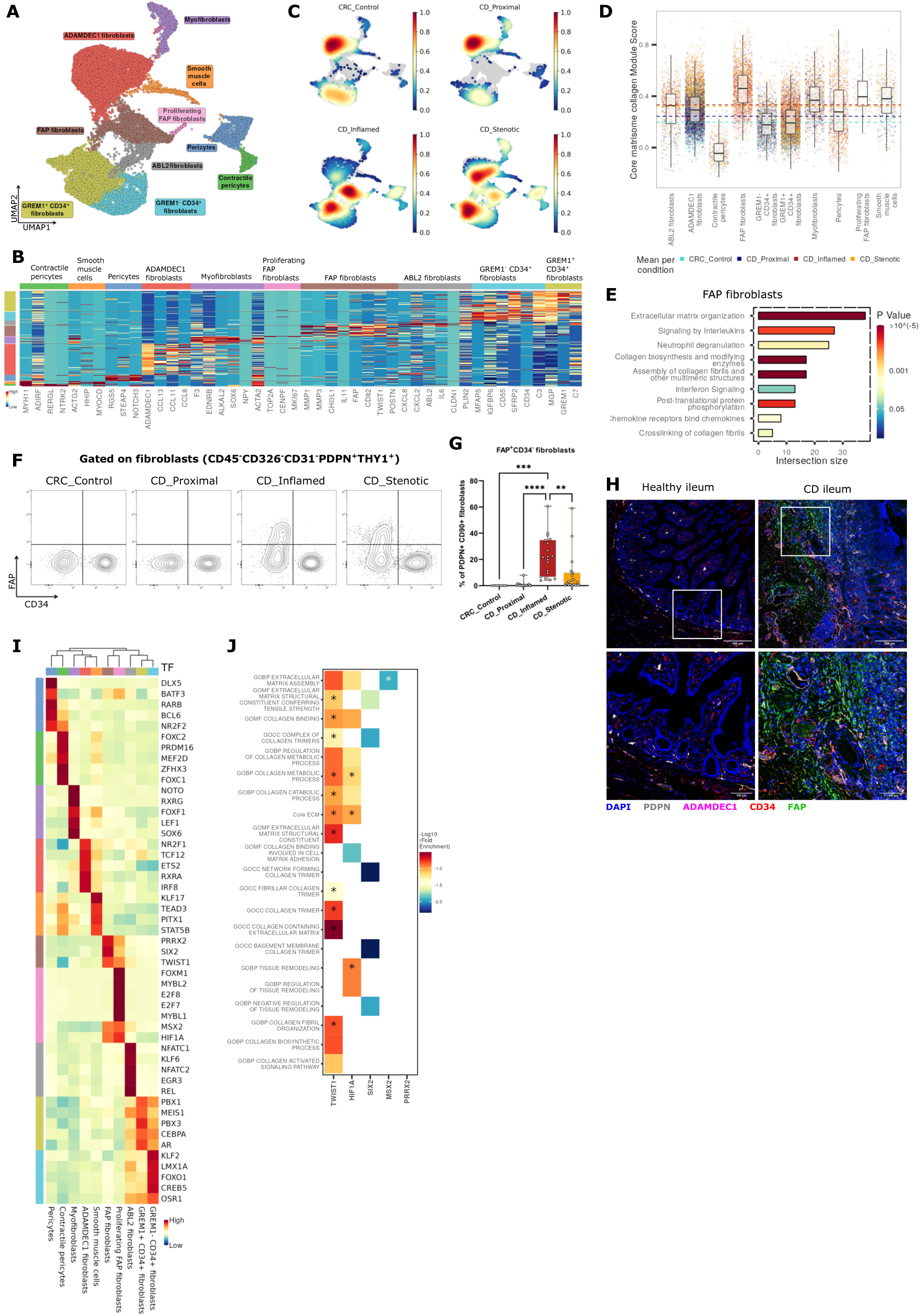
Heterogeneity of stromal cells in fibro-stenotic CD. (**A**) UMAP representation of re-clustered mesenchymal cells across different lesions of the terminal ileum. (**B**) Heatmap showing relative expression of top marker genes in each subset. (**C**) Cell subset composition across different lesions of the terminal ileum. (**D**) Bar plot showing gene set module score for core matrisome collagen genes in each stromal cell subset in different lesions. Horizontal lines indicate medians of respective lesions(**E**) Enrichment analysis for Reactome biological pathways in FAP fibroblasts (logFC, >0.5; FDR, <0.1). (**F**) Flow cytometry gating strategy for fibroblast subsets and (**G**) the plot of FAP expression in pan fibroblasts (7-AAD^-^CD45^-^CD31^-^ CD326^-^PDPN^+^THY1^+^) in different lesions of terminal ileum. Data are shown as box and whisker plots. Statistically significant differences were determined using a one-way ANOVA test corrected with Tukey’s multiple comparisons test (\*\**p* <0.01, \*\*\**p* <0.005, \*\*\*\**p* <0.001). (**H**) Immunofluorescence staining for PDPN, ADAMDEC1, CD34 and FAP expression in healthy ileum and CD diseased ileum. Original image composed of stitched 25× images. (**I**) Heatmap showing relative transcription factor activity in each stromal cell subset based on Single-cell regulatory network inference and clustering (SCENIC) analysis. (**J**) Heatmap showing selected terms after functional enrichment analysis of top 5 regulons using GO terms and core ECM gene set from MatrisomeDB (*indicate statistically significant terms after one sided Fisher’s exact test and multiple correction by Benjamini & Hochberg method).

Next, to confirm the presence of FAP fibroblasts in inflamed and stenotic samples, we performed flow cytometry on transmural ileum of CD patients. After gating out leukocytes (CD45), endothelial cells (CD31) and epithelial cells (CD326), we used CD90 and podoplanin (PDPN) to identify fibroblasts (Figure 2F and S2B). Using this approach, we confirmed that FAP fibroblasts (FAP^+^CD34^-^) were unique to the inflamed and stenotic ileum and absent in the control and proximal ileum (Figure 2G, S2 and S2D). In line, FAP enzymatic activity was significantly elevated in the inflamed and stenotic CD ileum compared to unaffected margins (Figure S2E).^19^ In addition to confirming the presence of FAP fibroblasts in inflamed and stenotic ileum of CD patients through flow cytometry, we also investigated the spatial distribution of the fibroblast subsets using multiplexed immunofluorescence staining (PDPN, CD34, FAP and ADAMDEC1). Importantly, we observed high expression of FAP in the submucosa and deeper layers of CD ileum where excess ECM deposition is observed. On the other hand, ADAMDEC1 fibroblasts were predominantly present in the healthy mucosa (Figure 2H and S2F).

FAP fibroblasts not only express higher ECM genes but are also characterised by an activated phenotype that over-expresses CD90, PDPN, and FAP proteins, pro-fibrotic autocrine loop molecules (*IL6, IL11, TGFB1*), neutrophils chemoattracting chemokines (*CXCL1, CXCL5, CXCL6*) and monocyte chemoattracting chemokines (*CCL2, CCL5, CCL7*) (Figure S2G). Moreover, single sample gene set enrichment analysis (ssGSEA) confirmed that FAP fibroblasts are enriched for processes such as inflammatory response and leukocyte chemotaxis (Figure S2H).^20^ Overall, these results suggested a key role for FAP fibroblasts in perpetual recruitment and potential activation of myeloid cells into the tissue during chronic inflammation.

In addition, we re-analysed and compared our data to previously published IBD scRNA-seq data sets of mucosal biopsies from CD and ulcerative colitis (UC) patients using Seurat integration and label transferring (Figure S6A).^14–16^ Cross-dataset cell type prediction score showed low to moderate similarity of FAP fibroblasts across these three datasets (Figure S2I). To further validate the presence of FAP fibroblasts across both CD and UC in colon, we analysed transmural samples from healthy CRC colon (unaffected), healthy CD colon (non-inflamed), inflamed/stenotic CD colon (granulating ulcer and thickened bowel wall) and UC colon (inflamed) using flow cytometry (Figure S2J). Of note, a significantly high proportion of FAP fibroblasts was exclusively noted in the inflamed/stenotic colon of patients with CD and not in those with inflamed UC colon (Figure S2K). These results suggest that FAP fibroblasts are a unique pathogenic subset present during chronic inflammation and fibro-stenosis, exhibiting excessive ECM deposition in CD.

### Transcriptional regulation and pro-inflammatory properties of FAP fibroblasts

Given that the unique transcriptional programs in FAP fibroblasts indicated distinct functions, we set out to investigate the upstream transcriptional regulators driving the unique transcriptional state in the FAP fibroblasts. Using single-cell regulatory network inference and clustering (SCENIC) analysis, TWIST1, SIX2, PRRX2, MSX2, and HIF1α were identified as the top regulons (a network of a transcription factor and corresponding targets) active in FAP fibroblasts (Figure 2I)^21^. To distinguish the main transcription factor driving the high ECM gene expression exhibited by FAP fibroblasts, we performed gene enrichment analysis of these top regulons. TWIST1 target genes stood out with significant enrichment of several GO terms associated with excessive ECM deposition such as collagen containing extracellular matrix as well as the gene set containing the core constituents of ECM (Figure 2K).

Consistent with the established role of TWIST1 in epithelial-mesenchymal transition (EMT), ssGSEA demonstrated a significant enrichment of EMT gene signatures in FAP fibroblasts (Figure S2L).^33, 34^ In addition, our ssGSEA analysis revealed a significant enrichment of terms associated with cellular senescence, including KEGG’s cellular senescence and Reactome’s senescence-associated secretory phenotype in FAP fibroblasts (Figure S2M) and enrichment of genes associated with the senescence-associated secretory phenotype (SASP) (Figure S2N).^22, 24, 26^ Overall, these results suggest that TWIST1 drives a pro-fibrotic phenotype in FAP fibroblast. In addition, the cellular senescence pathway may be active in FAP fibroblasts and could be driving the secretion of inflammatory cytokines and chemokines to worsen the inflammation on top of their pro-fibrotic function.

### GREM1^-^CD34^+^ fibroblasts as potential precursors of FAP fibroblasts

To gain insight into how FAP fibroblasts appear during chronic inflammation, we exploited the gene expression data to construct a differentiation trajectory for FAP fibroblasts. After identifying three fibroblast clusters topologically connected to FAP fibroblasts using partition-based graph abstraction (PAGA), we applied monocle3 on these four clusters (Figure S3A).^23, 25^ Trajectory analysis revealed that GREM1^-^CD34^+^ fibroblasts differentiated into FAP fibroblasts during inflammation (Figure 3A). Along the differentiation trajectory towards FAP fibroblasts, fibroblasts exhibited a loss of *CD34* expression and an acquisition of *TWIST1, FAP,* and *COL1A1* expression (Figure 3B and S3B). We hypothesised that the markedly distinct inflammatory microenvironment in CD may induce FAP fibroblasts differentiation from GREM1^-^CD34^+^ fibroblasts. To identify the specific factors that drive this differentiation, we used CellPhoneDB, a computational tool that estimates intercellular interactions between cell types and fibroblasts based on a curated database of ligand-receptor interactions.^27^ As expected, we observed an intense crosstalk within the mesenchymal compartment and between mesenchymal and endothelial cells. Beyond, myeloid cells were responsible for most of the predicted interactions with mesenchymal cells in general and particularly with FAP fibroblasts as lesions progressed to stenosis (Figure 3C and S3C).

**Figure 3.**
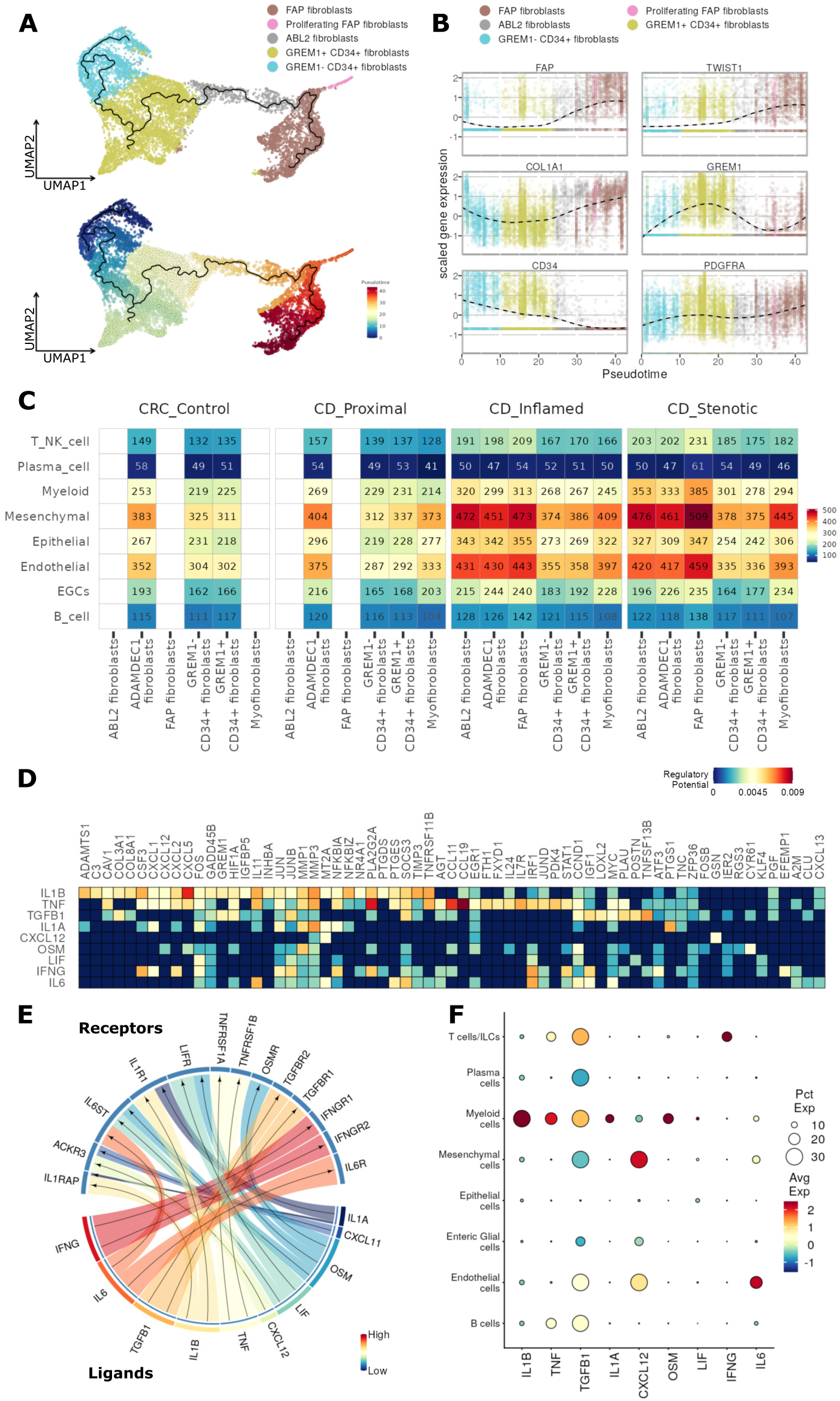
Trajectory analysis of fibroblast subset and stromal-immune interactions. (**A**) Pseudo-time trajectory projected onto a UMAP of selected fibroblast subsets. (**B**) Normalised expression levels of selected markers visualised along the pseudo-time. (**C**) Heatmap showing number of interactions (Ligand-Receptor pairs) between cell compartments and mesenchymal subsets. (**D**) Niche signalling driving FAP fibroblast differentiation, predicted by NicheNet; Regulatory potential of each target gene in columns by ligands in rows. (**E**) Circos plot depicting links between predicted ligands by NicheNet and their receptors. (**F**) Dot

### Stromal-immune cell interaction induce FAP fibroblast differentiation and activation

To better understand the signalling cues inducing the transcriptional shift resulting in the differentiation of FAP fibroblasts in CD ileum, we employed a recently described computational tool called NicheNet, which enables the prioritisation of ligands responsible for inducing alterations in a specific gene set.^28^ Based on the differentially regulated genes during the differentiation of FAP fibroblasts, NicheNet predicted *IL1A, IL1B, IL6, CXCL12, IFNG, LIF, OSM*, *TNF,* and *TGFB1* as the top ligands associated with the transcriptional reprogramming of GREM1^-^CD34^+^ fibroblasts towards FAP fibroblasts (Figure 3D and 3E). The myeloid compartment exhibited the highest gene expression levels of the highest ranked ligands including *IL1A*, *IL1B*, *TNF*, *TGFB1*, and *OSM*. In contrast, *IFNG* expression was predominantly found in T cells/ILCs, while *IL6* and *CXCL12* were mainly expressed by mesenchymal and endothelial cells (Figure 3F, and S3C). The activation of these signalling pathways was also confirmed by ssGSEA analysis (Figure S3D). Taken together, NicheNet analysis suggested that myeloid cells may be responsible for the *pro-fibrotic cues* responsible for FAP fibroblast differentiation and activation through several inflammatory mediators.

### scRNA-seq analysis reveals presence of a pro-fibrotic monocyte subset specific to inflammation and stenosis

To define in detail the pro-fibrotic myeloid cell subsets involved in driving the activation of FAP fibroblasts, we performed unsupervised re-clustering of the myeloid compartment (*LYZ, CTSG, CD68, CSF1R*) and we identified a total of fourteen clusters in the CD ileum and in the healthy CRC control ileum (Figure 4A, 4B and S4A). Two clusters of mast cells (*KIT, CTSG*) were identified in all segments, with minimal differences in gene expression observed between the two clusters (data not shown). Four distinct dendritic cell (DC) clusters were also identified including cDC1 (*CCND1, CLEC9A*), cDC2 (*LTB*), lymphoid DCs (*CCR7, LAMP3*) which were enriched in inflammation and plasmacytoid DCs (pDCs) which initially clustered along with B cells (*GZMB, TCF4, IRF7*) (Figure 4C). The macrophage and monocyte clusters exhibited remarkable variations across the different tissue conditions. Two macrophage clusters, distinguished by *LYVE1* or *IGF1* expression, were mainly present in non-inflamed ileum as opposed to inflamed and stenotic ileum. These clusters also displayed other typical mature resident macrophage markers such as *C1QA* and *C1QB* but lacked *CCR2* expression. Three clusters, namely CCR2 monocytes, MMP9 macrophages, and neutrophils were predominant in inflamed and stenotic ileum (Figure S4A). We also observed an inflammatory monocyte cluster (Inflammatory monocytes), which could be identified by the expression of *SLAMF1* (CD150) and common monocyte markers, including *CD300E*, *FCN1* and *CCR2*. Most strikingly, this cell subset was exclusively present in inflamed and stenotic ileum (Figure 4B and S4A). To identify the subset(s) of myeloid cells inducing FAP fibroblast differentiation during chronic inflammation, we evaluated the inflammation specific appearance and secretion of myeloid-derived pro-fibrotic ligands. Most importantly, Inflammatory monocytes had the highest expression of the pro-fibrotic ligands (*IL1A, IL1B, TNF*, and *TGFB1*) predicted by the NicheNet analysis (Figure 4D).

**Figure 4.**
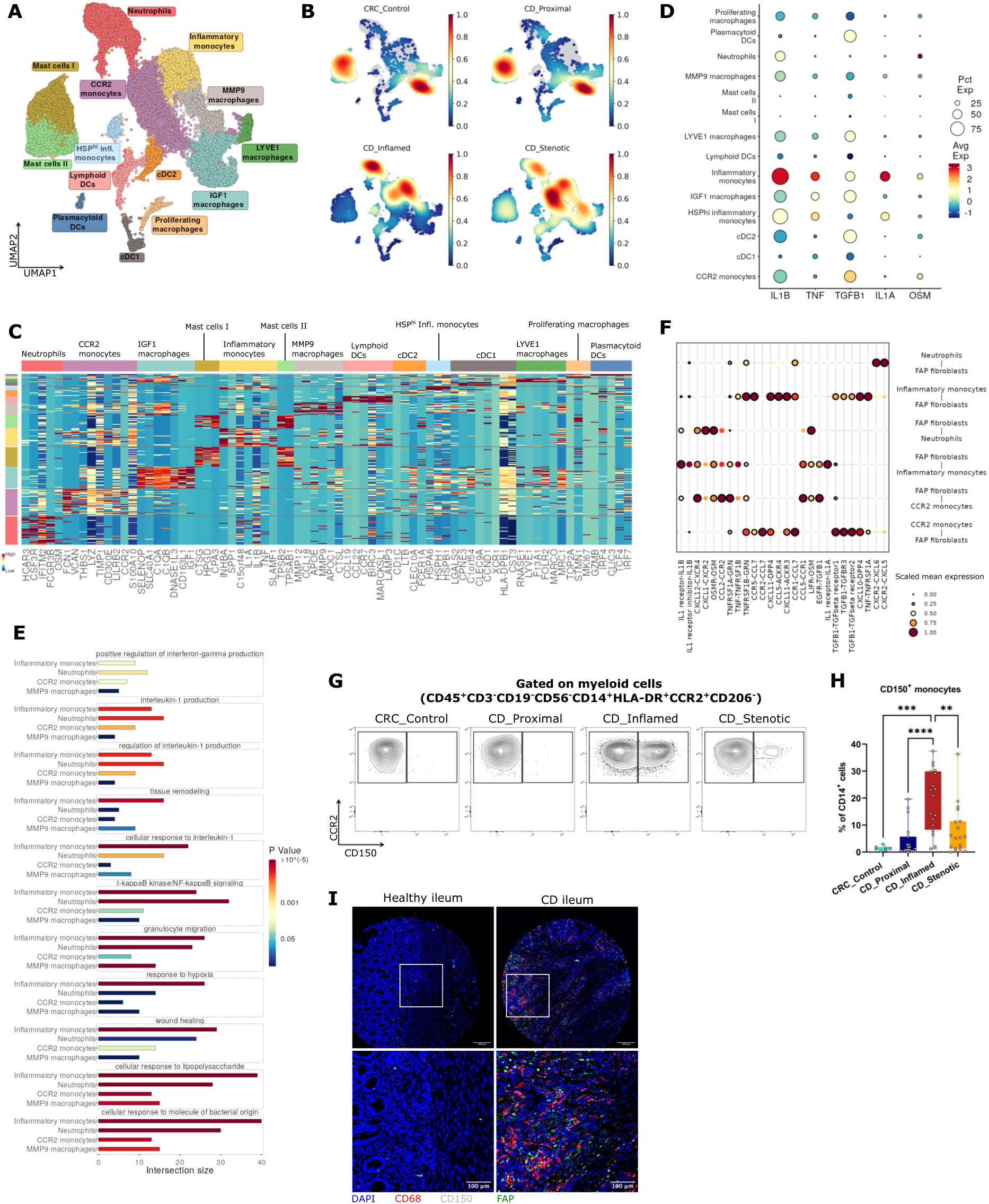
Heterogeneity of myeloid cells in fibro-stenotic CD. (**A)** UMAP representation of re-clustered myeloid cells and (**B**) cell subset composition across different lesions of the terminal ileum. (**C**) Heatmap showing the expression of the top marker genes of each myeloid subset. (**D**) Dotplot showing NicheNet predicted ligands expressed by myeloid cell subsets. (**E**) Selected Gene Ontology terms significantly enriched myeloid cell subsets. (**F**) Cellphone DB dot plot showing ligand-receptor interactions between FAP fibroblasts and Inflammatory monocytes or Neutrophils. First and second interacting molecules correspond to first and second cell types on the y axis respectively. Black circles indicate significant interactions (**G**) Flow cytometry gating strategy for myeloid cell sub-populations and (**H**) the plot of CD150 (*SLAMF1*) expression in CD14^+^ myeloid cells (7-AAD^-^CD45^+^CD3^-^CD19^-^CD56^-^HLA-DR^+/-^) in different lesions of terminal ileum. Data are shown as box and whisker plots. Statistically significant differences were determined using a one-way ANOVA test corrected with Tukey’s multiple comparisons test (\*\**p* <0.01, \*\*\**p* <0.005, \*\*\*\**p* <0.001). (**I**) Immunofluorescence staining for CD68, CD150 and FAP expression in healthy ileum and CD diseased ileum. Original image composed of stitched 25× images.

### Inflammatory monocytes as a hyper-inflammatory activation state of monocytes

Given the potential role of inflammatory monocytes in fibrosis, we further characterised this cluster. We compared the four myeloid clusters (Neutrophils, CCR2 monocytes, Inflammatory monocytes and MMP9 macrophages), which were predominantly present in inflamed and stenotic ileum. The functional gene enrichment analysis employing GO terms revealed a significant enrichment in Inflammatory monocytes, characterised by a substantial overlap between differentially up-regulated genes and GO gene sets such as (regulation of) interleukin-1 production, tissue remodelling, and wound healing, indicating their high inflammatory and pro-fibrotic profile compared to the other three myeloid subsets (Figure 4E). Similarly, Reactome pathway analysis revealed enrichment of pathways associated with extracellular matrix organisation, chemokine receptors bind chemokines, and interleukin-1 family signalling in Inflammatory monocytes. This underscores the inflammatory and pro-fibrotic characteristics of these cells (Figure S4B). Furthermore, we quantified the intercellular interactions between Inflammatory monocytes and other cell types using CellPhoneDB. Our analysis revealed that FAP fibroblasts exhibited a higher number of ligand-receptor interactions with Inflammatory monocytes in comparison to other immune cell subsets (Figure S4C). Of note, this analysis indicated a potential role for FAP fibroblasts in the recruitment of monocytes via CCL2, CCL5 and CCL7 (Figure S2G and 4I).

Further, to distinguish the main transcription factors driving the pro-fibrotic phenotype of Inflammatory monocytes, we performed SCENIC analysis. ELK3, MSC, STAT4, HIF1α, and IRF7 were found to be among the top significantly enriched regulons in Inflammatory monocytes (Figure S4D). Given the unique appearance of Inflammatory monocytes during inflammation, we hypothesised that they originate from monocytes during chronic inflammation. To understand the monocyte dynamics across disease states, we performed trajectory analysis of clusters connected with monocytes as identified by PAGA (data not shown). Nevertheless, we removed pDCs from the trajectory analysis as they have been reported to be not of monocyte origin.^31^ Trajectory analysis showed two branches of monocyte differentiation through pseudotime (Figure S4E and S4F). One branch of CCR2 monocytes gave rise to IGF1 macrophages, with a subsidiary branch giving rise to cDC2. This branch was predominant in control and proximal ileum. The other branch depicted CCR2 monocytes differentiating into Inflammatory monocytes and further to MMP9 macrophages predominantly in inflamed and stenotic ileum (Figure 4B, S4E and S4F).

### Inflammatory monocytes are predominantly present in inflamed and stenotic ileum

To further confirm the association of the Inflammatory monocytes with inflammation and stenosis in transmural CD ileum, we performed flow cytometry and immunofluorescence staining, using CD150 as a reliable marker to identify Inflammatory monocytes within the myeloid compartment. In the diseased ileum, CD14^dim^ cells (as Neutrophils), CD150^-^ monocytes (as CCR2 monocytes) and CD150^+^ monocytes (as Inflammatory monocytes) were significantly increased, compared to the control and proximal ileum (*p* < 0.005) (Figure 4G, 4H, S4G and S4H). Similarly, with multiplex immunofluorescent staining, we observed an increase of CD68^+^CD150^+^ cells (as Inflammatory monocytes) in the deeper muscularis layer of diseased CD ileum, which co-localised with FAP expression (Figure 4I and S4I). We confirmed CD150 as a surface marker for Inflammatory monocytes within the myeloid compartment, facilitating their isolation for *in vitro* experiments. Taken together, Inflammatory monocytes are a pathogenic subset of myeloid cells that secrete an excess of inflammatory and pro-fibrotic mediators, inducing differentiation and activation of FAP fibroblasts leading to tissue remodelling.

### CD150 Inflammatory monocytes are a key feature of CD

We identified Inflammatory monocytes in our scRNA-seq analysis of transmural CD ileum and confirmed them by flow cytometry. To contextualise our findings within the broader research landscape, we re-analysed and compared our data to three IBD scRNA-seq data sets derived from mucosal biopsies of both CD and UC patients using Seurat integration and label transferring (Figure S6B). Similar myeloid clusters were identified in all three IBD datasets with low to moderate similarity score (Figure S4J). To further confirm the presence of Inflammatory monocytes across IBD, we performed flow cytometry analysis by using transmural colon from CD and UC patients (Figure S4K). Notably, the presence of CD150^+^ monocyte population was unique to CD in the colon (Figure S4K). Together with our findings in CD ileum, these results suggest that CD150 Inflammatory monocytes are a unique subset with highly inflammatory and pro-fibrotic properties, specific to CD patients in both inflamed ileum and colon.

### Inflammatory monocyte derived *pro-fibrotic cues* activate FAP fibroblasts via TWIST1

To validate the expression and functional relevance of pro-fibrotic ligands in Inflammatory monocytes as observed in our scRNA-seq data, we quantified the gene expression of pro-fibrotic ligands across different FACS-sorted myeloid subsets: HLA-DR^+^ cells, CD14^dim^ neutrophils (CD14^low^, HLA-DR^neg^), CD150^-^ monocytes and CD150^+^ monocytes (Figure S5A). In line with our scRNA-seq data, Inflammatory monocytes (CD150^+^ monocytes) showed the highest expression of *IL1A, IL1B, TNF, TGFB1*, and *SLAMF1* but not *OSM*, which was expressed mainly by the CD14^dim^ neutrophils as previously reported (Figure 5A).^56^ Next, we sought to validate the capacity of Inflammatory monocytes to induce fibroblast activation. To this end, we performed *in vitro* fibroblast activation assays with CCD-18Co fibroblasts (CRL-1459, ATCC) and primary ileal fibroblasts isolated from transmural ileum of healthy regions from CRC patients. To confirm that Inflammatory monocyte-derived ligands modulate fibroblast activation as seen in our scRNA-seq data, we sorted CD14^dim^ neutrophils, CD150^-^ monocytes and CD150^+^ monocytes and collected cell supernatant after 16 hours of culture. As shown in our computational analysis, the supernatants of Inflammatory monocytes isolated from inflamed ileum (5 CD patients undergoing surgery for ileal stenosis) induced an activated fibroblast state with up-regulated expression of TWIST1, FAP and type III collagen compared to the supernatants of other FACS-sorted myeloid subsets (Figure 5B, 5C and S5B). To further confirm that ECM deposition induced by Inflammatory monocyte-derived ligands modulate fibroblasts, we co-cultured fibroblasts and FACS-sorted myeloid cell supernatants with ascorbic acid for three weeks and performed immunofluorescent staining without permeabilisation. Cell supernatants of Inflammatory monocytes promoted significantly higher secreted FAP expression and higher extracellular deposition of type I collagen, and type III collagen by fibroblasts, compared to the control and CD14^dim^ neutrophils (Figure 5D, 5E and S5C).

**Figure 5.**
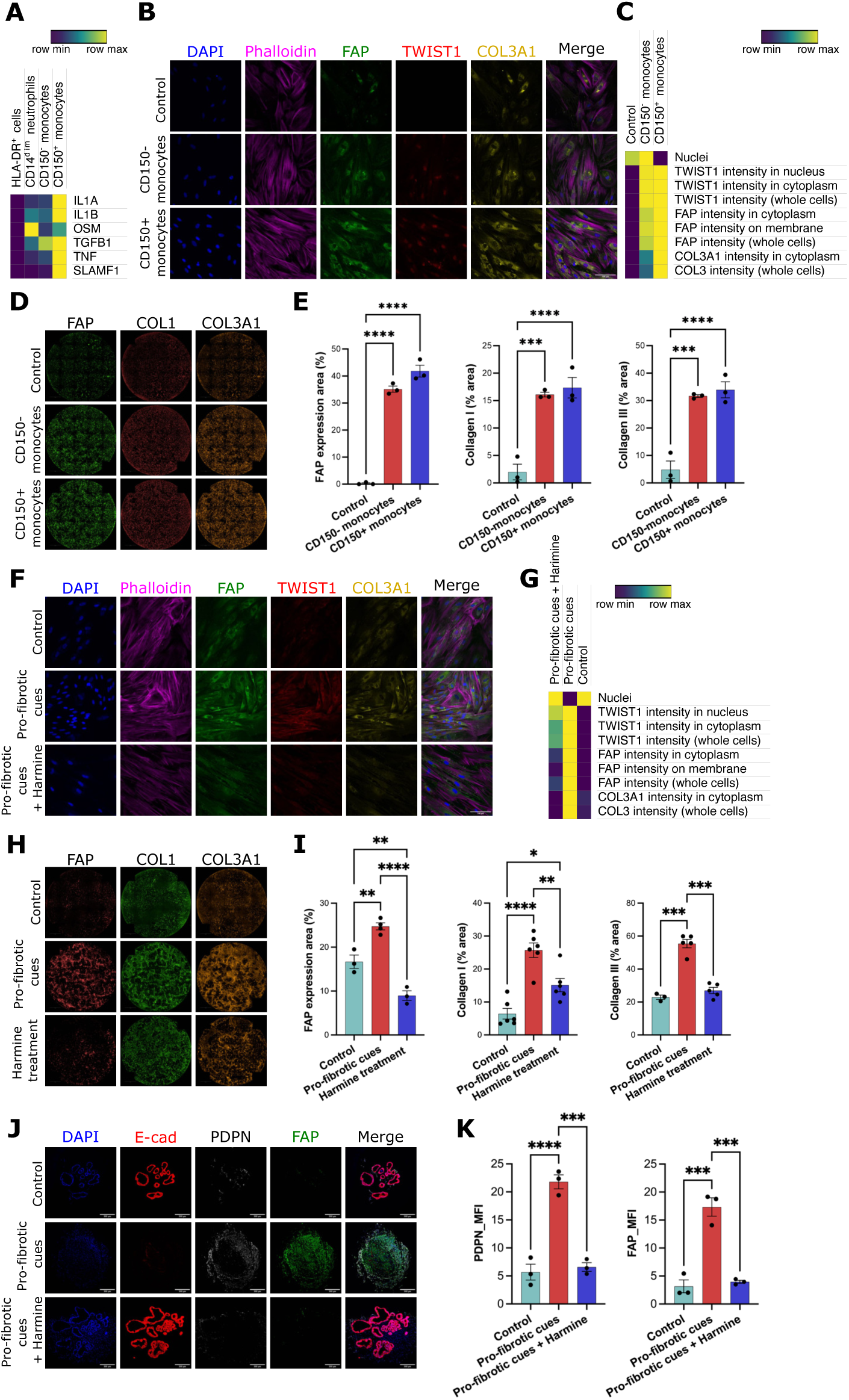
CD150^+^ monocytes-derived cytokines promoted FAP fibroblast activation and extra-cellular matrix protein deposition. (**A**) Heatmap showing relative expression of NicheNet-predicted ligands expressed by FACS-sorted myeloid cell subsets (n=4). (**B**) Immunofluorescence staining (25× image) and (**C**) Heatmap showing relative expression of FAP, TWIST1 and type III collagen in monocyte-stimulated CCD-18Co fibroblasts. (**D**) Immunofluorescence staining (10× image) and (**E**) Bar plot showing relative expression of FAP, type I and type III collagen in monocyte-stimulated CCD-18Co fibroblasts. Data are shown as bar plots with SEM. Statistically significant differences were determined using a one-way ANOVA test corrected with Tukey’s multiple comparisons test (\*\*\**p* <0.005, \*\*\*\**p* <0.001). (**F**) Immunofluorescence staining (25× image) and (**G**) Heatmap showing relative expression of FAP, TWIST1 and type III collagen in *pro-fibrotic cues*-stimulated CCD-18Co fibroblasts. (**H**) Immunofluorescence staining (10× image) and (**I**) Bar plot showing relative expression of FAP, type I and type III collagen in *pro-fibrotic cues*-stimulated CCD-18Co fibroblasts after TWIST1 inhibition. Data are shown as bar plot with SEM. (**J**) Immunofluorescence staining (10x image) and (**K**) quantitative analysis (mean fluorescence intensity) of FAP and PDPN *pro-fibrotic cues*-stimulated iPSC-derived intestinal organoids with or without Harmine. Data are shown as bar plot with SEM. Statistically significant differences were determined using a one-way ANOVA test corrected with Tukey’s multiple comparisons test (\**p* <0.05, \*\**p* <0.01, \*\*\**p* <0.005, \*\*\*\**p* <0.001).

Next, to assess the potency of different cytokine combinations in inducing fibroblast activation, we monitored response of fibroblasts to different cytokine combinations using high-content imaging. The combination of IL-1α, IL-1β, TNFα, TGFβ1, OSM and IFNγ (henceforth referred to as *pro-fibrotic cues*) induced the highest expression of TWIST1, FAP and type III collagen among different cytokine combinations after 48 hours of stimulation (Figure S5D). Computational analysis had indicated a specific transcriptional regulatory role for TWIST1 in the excess ECM deposition exhibited by FAP fibroblasts. Hence, we investigated if inhibition of TWIST1 could affect the pro-fibrotic nature of FAP fibroblasts. To confirm the involvement of TWIST1 in fibroblast activation and ECM deposition, we used Harmine, a recently identified TWIST1 inhibitor, that induces the degradation of TWIST1.^39, 41^ After Harmine treatment, *pro-fibrotic cues*-stimulated fibroblasts showed a remarkable reduction in the expression of FAP, TWIST1 and type III collagen compared to vehicle treated cells (Figure 5F and 5G). Next, we further determined the effect of *pro-fibrotic cues* on ECM deposition *in vitro* (Figure S5E). Extracellular staining showed that *pro-fibrotic cues* significantly promoted ECM deposition by primary fibroblasts, including fibronectin, type I, type III and type IV collagen. Notably, therapeutic intervention with Harmine effectively attenuated ECM deposition *in vitro* even after activation of the fibroblast (Figure 5H and 5I, S5F and S5G). Additionally, using a 3D *in vitro* model based on human iPSC-derived intestinal organoids (IOs) typically containing epithelium and stromal cells, we showed that the *pro-fibrotic cues* induced a significant increase of FAP and PDPN expression which was significantly reduced in the presence of Harmine (Figure 5J and 5K). Altogether, these results provided solid evidence that Inflammatory monocyte-derived *pro-fibrotic cues* modulates FAP fibroblast activation and ECM secretion in a TWIST1 dependent manner.

### Inhibition of TWIST1 attenuates chronic DSS colitis-induced intestinal fibrosis

Based on our scRNA-seq data analysis and *in vitro* experiments, we identified TWIST1 as a critical transcription factor involved in fibroblast activation and intestinal fibrosis. To corroborate this observation *in vivo*, we used a chronic DSS colitis-induced fibrosis model to examine the involvement of TWIST1 in ECM deposition and fibrotic processes. The disease activity index (DAI) analysis showed that Harmine-treated mice had an improved recovery during the first and second cycles of DSS treatment but not in the third cycle compared to the DSS-only group (Figure S6C to S6F). Despite the absence of a significant difference in the DAI, Harmine treatment led to reduced collagen deposition in DSS-treated mice (*p* < 0.05) (Figure 6A and 6B), along with a decrease in TWIST1 and FAP expression as demonstrated by immunofluorescent staining (Figure 6C). These results demonstrated that inhibition of TWIST1 could be a potential therapeutic strategy to attenuate intestinal fibrosis.

**Figure 6.**
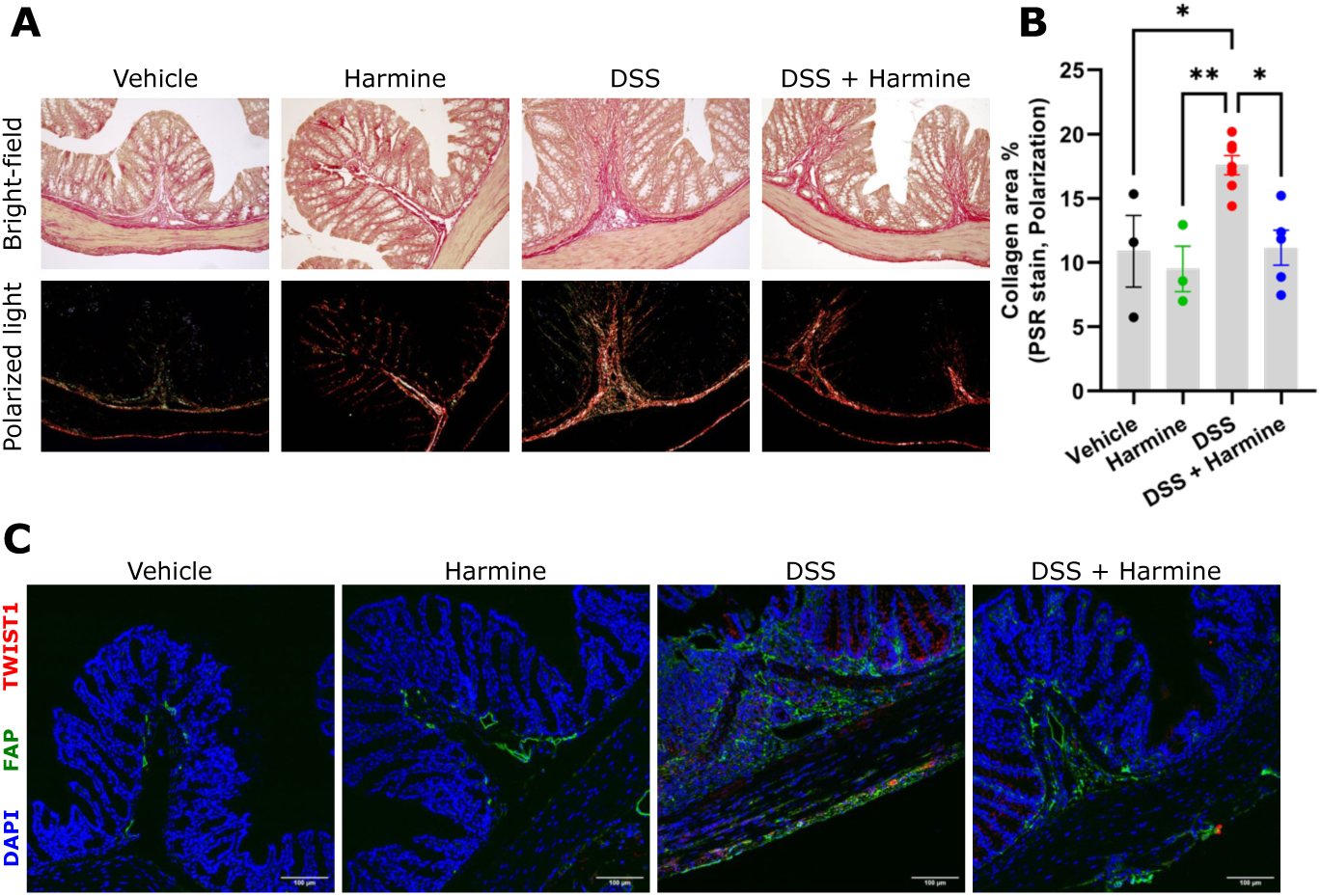
TWIST1 inhibition attenuated intestinal fibrosis. (**A**) Picrosirius red staining and (**B)** Bar plot showing collagen deposition in mouse colon of chronic DSS colitis after Harmine treatment. Data are shown as bar plots with SEM. Statistically significant differences were determined using a one-way ANOVA test corrected with Tukey’s multiple comparisons test (\**p* <0.05, \*\**p* <0.01). (**C**) Immunofluorescence staining for FAP and TWIST1 expression in mouse colon of chronic DSS colitis after Harmine treatment. Original image composed of stitched 25× images.

## DISCUSSION

Lack of transmural sampling has been a long-standing limitation in CD studies despite the disease occurring in all the layer of the gut wall.^29^ So far, scRNA-seq studies in CD have mostly focused on characterising mucosal inflammation using endoscopic biopsies and hence do not appreciate the transmural heterogeneity in CD ileum.^14–16^ Our study is the first of its kind to utilise transmural ileal biopsies for scRNA-seq, allowing the cellular profiling of CD lesions in the deeper layers of the gut, where most of the tissue remodelling is detected in patients.

Analysis of our scRNA-seq data revealed a previously unknown heterogeneity in the transmural CD ileum. Overall, fibroblast and myeloid compartments showed remarkable differences across disease conditions. Deeper analysis of the mesenchymal compartment revealed FAP fibroblasts as the key pathogenic cell subset uniquely present in the inflamed and stenotic CD ileum, responsible for excessive deposition of ECM. Importantly, we identified a crucial effector role for a distinct subset of inflammatory monocytes in activating FAP fibroblasts via inflammatory mediators. Reinforcing this observation, we could spatially locate FAP fibroblasts in the deeper submucosa and muscularis layers of fibro-stenotic CD ileum in close proximity with inflammatory monocytes expressing SLAMF1 (CD150), emphasizing their potential interaction in the pathological process.

Notably, FAP fibroblasts exhibited elevated expression of collagen and ECM genes, along with an activated phenotype characterised by over-expression of pro-fibrotic autocrine loop molecules, such as *IL11* and *IL6*, and chemokines for neutrophils (*CXCL1*, *CXCL5*, *CXCL6*) and macrophages (*CCL2*, *CCL5*, *CCL7*), suggesting a critical role in perpetuating a feed-forward loop that sustains chronic inflammation within the tissue. Trajectory analysis indicates that FAP fibroblasts differentiate from homeostatic CD34^+^GREM1^-^ fibroblasts during inflammation, which aligns with prior studies demonstrating that CD34 mesenchymal cells are essential for maintaining intestinal homeostasis and differentiate into myofibroblasts via the TGFb-Smad2 signalling pathway in response to inflammation.^30^ Similar observation has been made in the heart, where depletion of CD34^+^ cells lead to reduced myocardial fibrosis and improved cardiac function.^32^

Transcriptional regulatory network analysis revealed TWIST1 as the main transcriptional regulator driving the excess ECM gene expression by FAP fibroblasts. TWIST1 has been primarily investigated in cancer for its role in EMT.^33, 34^ Concordantly, our gene enrichment analysis indicated an activation of EMT in FAP fibroblasts. In line with our findings, TWIST1 has also been implicated in other fibrotic diseases such as pulmonary fibrosis and renal fibrosis.^35^ Moreover, TWIST1 has demonstrated pro-fibrotic characteristics in human fibroblasts by enhancing matrix stiffness.^36^ We confirmed the role of TWIST1 in the activation of FAP fibroblasts using Harmine. Harmine, as a TWIST1 inhibitor, has been used *in vitro* and *in vivo* studies to target EMT, tumorigenesis and cancer cell invasion.^39, 41^ In our study, Harmine treatment attenuated *pro-fibrotic cues*-induced fibroblast activation and ECM protein secretion and deposition *in vitro*. Additionally, i*n vivo* experiments demonstrated that inhibition of TWIST1 by administration of Harmine attenuated ECM deposition in mice subjected to chronic DSS-colitis, further corroborating the significance of TWIST1 in the fibrotic process. In line with the proposed role for FAP fibroblast in leukocyte chemotaxis and secretion of pro-inflammatory cytokines, a previous study found that TWIST1 inhibition by Harmine reduced leukocyte infiltration and secretion of pro-inflammatory cytokines in wild-type mice with DSS-hypoxia-induced colitis.^37^

To further elucidate the environmental cues driving differentiation of activated fibroblasts, we employed computational analysis using intercellular communication tools. In addition to autocrine communication within the stromal compartment, ligand-receptor pair analysis revealed extensive crosstalk between myeloid cells and FAP fibroblasts, particularly during inflammation and stenosis. The pro-fibrotic microenvironmental cues were specifically composed of myeloid-derived cytokines, including IL1β, IL1α, OSM, and TNFα, which induced the differentiation of FAP fibroblasts. While monocyte accumulation is a crucial aspect of CD, the precise effector role of monocytes in the progression of severe disease and dysregulated tissue remodelling remains poorly understood.^44^ Monocytes exhibit remarkable plasticity in adapting their functional state to the changes in the microenvironment, enabling them to guide stromal cells either towards regulated wound healing, or towards dysregulated tissue remodeling.^38, 40^ Our study clearly indicate that chronic inflammatory state drives myeloid cells in a hyperinflammatory state resulting into dysregulated tissue remodelling by the activation of pathological fibroblast differentiation.

Deeper characterization of the myeloid cells revealed that the pro-fibrotic signals were specifically secreted by a subset of monocytes identified by the unique expression of *SLAMF1* (CD150). These inflammatory myeloid cells differentiated from monocytes during inflammation and stenosis presented with a hyper-inflammatory signature with high expression of inflammatory mediators. Further substantiating their activated phenotype, the transcriptional regulation in these monocytes were distinctly driven by the transcription factors known for inflammatory monocyte state such as STAT4, HIF1α, MSC among others.^42, 43, 45^ Of note, CD150 Inflammatory monocytes co-localised with FAP fibroblast in the deeper submucosa and muscularis layer in inflamed and stenotic CD lesions. Our observations are in line with a recent investigation, which demonstrated that when stimulated with lipopolysaccharide (LPS), blood-derived monocytes from CD patients resistant to anti-TNF therapy displayed a hyper-inflammatory phenotype.^46^ This was characterised by an increased release of inflammatory cytokines, such as TNFα, IL-23, and IL-1β. Taken together, these discoveries imply a potential association between a dysregulated immune response in monocytes, the development of fibrosis, and resistance to anti-TNF therapy in certain CD patients.

Our findings are consistent with the emerging paradigm of the crucial role for stromal cells in chronic inflammation. A pathological stromal cell subset of FAP fibroblasts with pro-fibrotic and pro-inflammatory characteristics has recently been observed in various organs including liver, lung, heart, joint and multiple cancers.^47, 49, 51, 53–55^ Depletion of FAP expressing fibroblasts using engineered FAP CAR-T cells, have proved to reduce tissue damage and fibrosis in experimental models of AngII/PE-induced cardiac fibrosis and in bleomycin-induced lung fibrosis.^48, 50, 52^ In addition, depletion of FAP expressing cells led to reduced leukocyte infiltration and severity in a murine model of arthritis.^51^ In CD as well, FAP fibroblasts and inflammatory macrophages have previously been reported in ileal lesions of IBD patients who have failed anti-TNF therapy.^14^ In line, recent study in CD has shown induction of FAP fibroblasts via neutrophil derived IL-1 signalling.^56^ However, neutrophils are mostly present at the luminal side of the mucosa surrounding ulcers. Hence, the neutrophil mediated activation of fibroblasts alone does not explain the presence of FAP fibroblasts in the deeper fibrotic ileum of CD patients as we have observed.

To contextualise our findings in the broader understanding of cellular heterogeneity in IBD, we compared Inflammatory monocyte and FAP fibroblast transcriptional signatures across the major IBD scRNA-seq data sets (Figure S6A and S6B).^14–16^ In Kong et al.’s data, signature scores for FAP fibroblasts and Inflammatory monocytes in ileum samples were consistently low, despite a considerable number of stricturing CD patients included in the study. In contrast, Martin et al. utilised resected mucosal samples and identified a distinct subset of anti-TNF therapy-resistant patients with high gene signature score for both cell types in the ileal lamina propria. This discrepancy may be attributed to Kong et al. ’s sampling strategy, which involved characterisation of endoscopic biopsy limited to the superficial mucosal layer. We postulate that the key difference between fibro-stenosis and chronic inflammation in patients with FAP fibroblasts may lie in the localisation of FAP fibroblasts in the deeper layers, rather than solely in the mucosa, as observed in anti-TNF therapy-resistant patients. Only a very minor subset of UC patients had both cell type signatures in the colon. In line, our flow cytometry analysis on transmural colon samples from CD and UC patients demonstrated that FAP fibroblast and CD150 Inflammatory monocytes are predominantly a feature of CD.

Overall, our study extends our knowledge on cellular heterogeneity in the transmural ileum of fibro-stenotic CD patients and highlights a novel intercellular interaction between immune cells and fibroblasts. We found that Inflammatory monocytes promote tissue remodelling and fibrosis by inducing the differentiation of a pro-fibrotic fibroblast subset during inflammation and stenosis. We also identified TWIST1 as a major transcriptional regulator of fibroblast activation leading to excessive ECM deposition. In conclusion, our study unravelled multiple potential therapeutic targets that hold significant promise for the development of new and more effective treatments for fibro-stenotic CD.

## Author contributions

Conceptualisation: BJK, SA, BV, GDH, SV, GM ; Methodology: BJK, GD, SA, GB, VDS, PVDV, IDM, JF, GDH, GM ; Software: SA, SS ; Formal analysis: BJK, SA ; Investigation: BJK, SA, GD, FB, SS, AZ, LVB, VB, YVR, EV, SaV, CB, BV, GDH ; Resources GB, AW, ADH, GDH, SA, BV, SV, GM ; Data Curation: BJK, SA, BV ; Visualisation: BJK, SA ; Funding acquisition: MS, SV, GM; Project administration: SV, GM; Supervision: SV, GM; Writing – original draft: BJK, SA; Writing – review & editing: all.

## Supporting information

Supplemental item Table S1

Supplemental item Table S2

Supplemental item Table S3

## Acknowledgments

We thank all patients for their participation in the study and for their willingness to donate tissue samples for research. The authors would like to thank Brecht Creyns (KU Leuven) for sample collection, Iris Appeltans, Karlien Vranken and Renata Siqueira de Mello (KU Leuven) for technical assistance, Sales Ibiza Martinez (UAntwerp) for suggestions on experimental setup, the IBD Leuven group for assistance with biobanking and administration, Reena Chinnaraj and Vera Dermesrobian (FACS Core, KU Leuven) for support with flow cytometry and sorting, Genomics Core (UZ Leuven) for assistance with scRNA-seq, Nikky Corthout and Axelle Kerstens BioImaging Core Leuven for assistance with the PerkinElmer Operetta CLS, and Nikon-Marzhauser Slide Express. Authors would like to thank Tobie Martens for his assistance and support with Zeiss LSM 780 – SP Mai Tai HP DS (Cell and Tissue Imaging Cluster, KU Leuven), supported by Hercules AKUL/11/37 and FWO G.0929.15 to Prof. Pieter Vanden Berghe. The computational resources were provided by the Flemish Supercomputer Center, funded by the Research Foundation - Flanders (FWO) and the Flemish Government. BioRender was used for making graphical images.

This study was supported by funding from Taiwan (Ministry of Education)-KU Leuven Scholarship Programme (BJK), Stichting tegen Kanker (VDS), FWO postdoctoral fellowship (SaV), KU Leuven Global PhD-University of Edinburgh Partnerships (GPUE/20, FB), KU Leuven Global PhD-University of Melbourne Partnerships (GPUM/22/020, SS), FWO grants (G086721N, GM and S008419N, SV and GM), FWO SB fellowship (1S64222N, YVR), KU Leuven Internal Funds (C12/15/016, GM and C14/17/097, GM, CB and SV), Clinical Research Fund (KOOR) at the University Hospitals Leuven (CB, BV), Research Council at the KU Leuven (BV), European Regional Development Fund-Project (CZ.02.1.01/0.0/0.0/16_019/0000868, VB and JF) and Ministry of Health of the Czech Republic-DRO (00023736, VB and JF).

## STAR Methods

### Human specimens

The resected terminal ileum was collected immediately after surgery from patients with Crohn’s disease (CD) or colorectal cancer (CRC) under the supervision of the specialised IBD-pathologist (GDH). The protocol was approved by the Institutional Review Board (IRB) of the University Hospitals Leuven, Belgium (B322201213950/S53684, CCARE, S-53684). All recruitment was performed after ethical approval and oversight from the IRB and informed consent was obtained from all participants before surgery. Clinical information and metadata for the samples in this study were provided in Supplemental item Table S3.

### Histological slides of the terminal ileum

Transmural biopsies from the terminal ileum were fixed in 4% formaldehyde, embedded in paraffin and 5µm thick sections were cut for histological analysis. Hematoxylin and eosin (H&E) and Masson’s trichrome staining were performed in the Department of Imaging & Pathology at the UZ Leuven. The pathological score system was modified from Gordon et al. (as Supplemental item Table S1 and S2) and examination was performed by a specialised IBD-pathologist (GDH).^17^ To quantify the relative histologic area of collagen on Masson’s trichrome stained slides, the average of ten images in each sample was taken and quantified with Image J. The slides were imaged on the Marzhauser Slide Express 2 (Nikon) at the VIB Leuven.

### Single-cell isolation from the terminal ileum

Single-cell suspensions were prepared from the transmural terminal ileum. Briefly, healthy (as the proximal tissue), granulating ulcerative (as the inflamed tissue) and thicken (as the stenotic tissue) biopsies of CD ileum and healthy (as the control group) ileum of CRC were treated with 1 mM DTT and 1 mM EDTA in 1x Hank’s balanced salt solution (HBSS), and 1 mM EDTA in HBSS at 37 °C for 30 minutes, respectively. Then the tissue was minced and digested with 5.4 U/mL collagenase D (Roche Applied Science), 100 U/mL DNase I (Sigma), and 39.6 U/mL dispase II (Gibco) in a sterile gentleMACS C tube for 20 minutes at 37 °C at 250 to 300 rpm after dissociating with the gentleMACS™ Dissociator (program human_tumor_02.01). After being treated with Red Blood Cell Lysis Buffer (11814389001, Roche), single-cell suspensions were used for scRNA-seq, cell culture and flow cytometry.

### FAP activity measurement

Snap-frozen ileum was crushed in a pre-cooled mortar kept on dry ice, intermittently using liquid nitrogen to keep samples frozen and to avoid loss of proteolytic activity due to temperature increase. Afterwards, the samples were lysed (1:5 ratio of sample (mg): lysis buffer (μL)) in lysis buffer (50 mM Tris-HCl pH 8.3, 10 mM EDTA, 1% n-octyl-β-D-glucoside and 70 μg/mL aprotinin) for 30 minutes on ice with frequent agitation. Next, samples were centrifuged at 12.000xg for 10 minutes at 4 °C. The ileum supernatant was collected and used immediately to perform a FAP enzymatic activity measurement and Bradford protein quantification assay. FAP enzymatic activity measurements were performed using the in-house developed assay using Z-Gly-Pro-AMC as the fluorogenic substrate and UAMC-1110 as specific FAP-inhibitor.^19, 57^

Briefly, in a 96-well plate (half-area, Greiner Bio-One), 5 µL ileum supernatant was pre-incubated for 15 min at 37 °C with 10 µL of FAP inhibitor or solvent control (250 nM UAMC-1110 or 0.0025% (v:v) DMSO in FAP assay buffer consisting of 100 mM Tris-HCl pH 8.0, 300 mM NaF, 1 mM EDTA and 50 mM salicylic acid). Afterward, 35 µL pre-heated substrate solution (Z-Gly-Pro-AMC in FAP assay buffer, final concentration 266 µM; Bachem, Bübendorf, Switzerland, cat nr: 4002518) was added and fluorescence was measured kinetically for 30 minutes at 37 °C using the Tecan Infinite® M200 Pro (excitation wavelength 380 nm and emission wavelength 465 nm; Tecan, Männedorf, Switzerland). Fluorescence intensity was related to an AMC standard curve (0.3125 μM-10 μM) in an identical buffer. FAP enzymatic activity was normalised to the total protein content in the samples by a Bradford protein quantification assay.

### Immunofluorescence staining of human ileum

The terminal ileum was fixed in 4% formaldehyde, embedded in paraffin and cut into 5µm thickness. Dewaxed and rehydrated 5 µm thick sections from the paraffin blocks s were subjected to heat-induced antigen retrieval by boiling in sodium citrate buffer (pH 6.0) for 20 minutes at 95°C (water bath). This was followed by 10 minutes of permeabilisation buffer (0.2% Triton X-100 in PBST) and 60 minutes of blocking in 2% BSA/PBST at room temperature. Tissue slides were labelled with primary antibodies in 2% BSA/PBST overnight at 4 °C, including sheep anti-human FAP antibody (1:40, AF3715, R&D Systems), mouse anti-human CD34 antibody (1:100, clone QBEnd/10, Sigma-Aldrich), rabbit anti-human ADAMDEC1 antibody (1:100, PA5-103574, Invitrogen), rabbit anti-human CD150 antibody (1:300, PA5-84809, Invitrogen), mouse anti-human CD68 antibody (1:100, clone KP1, BioLegend), and rat anti-human podoplanin antibody (1:300, clone NZ-1.3, eBioscience). Following incubation, the tissue slides were labelled with secondary antibodies in 2 % BSA/PBST for 2 hours at room temperature, including Alexa Fluor 488-conjugated donkey anti-sheep IgG antibody (A-11015, Invitrogen), Cy3-conjugated donkey anti-goat IgG antibody (705-165-003, Jackson ImmunoResearch), Cy3-conjugated donkey anti-rat IgG antibody (712-165-150, Jackson ImmunoResearch), Cy5-conjugated donkey anti-mouse IgG antibody (715-175-151, Jackson ImmunoResearch) and ATTO490LS-conjugated goat anti-rabbit IgG antibody (2309-1MG, Hypermol).

### Single-cell-RNA-sequencing and data analysis

Cell suspensions were processed with a 10x3’ v3 GEM kit and loaded on a 10x chromium controller to create Single Cell Gel beads in Emulsion (GEM). A cDNA library was created using a 10x 3’ v3 library kit and was then sequenced on a NovaSeq 6000 system (Illumina). Pre-processing of the samples including alignment and counting was performed using Cell Ranger Software from 10x.

Doublet score was calculated using three methods (scDblfinder, scrublet and DoubletFinder) on each sample separately and corresponding doublet scores were added to the metadata.^58, 60, 62^ Also using the DropletQC package, a QC metric indicating the fraction of reads exclusive to nuclear reads was calculated.^64^ Thus, our QC metrics included fraction of nuclear reads, doublet scores and nUMI.

Data were analysed using Seurat v3 SCTransform-Integration workflow with each patient as a batch. Only cells with more than 199 unique genes and less than 30 % mitochondrial genes were included in the analysis. Next, SCTransfrom function from Seurat was used to scale, normalise and transform each sample (10x channel) with method set as ‘glmgGamPoi’ and percentage of mitochondrial genes as the variable to regress.^59, 61, 63^ Next, 3000 features were selected for integration and a reference-based integration with 10 out of the 35 samples as reference samples was performed. After PCA, 80 principal components were used to find shared nearest neighbours and compute UMAP. Next, the shared nearest neighbour graph was used for clustering the cells at a resolution of 2.

After clustering, 2 clusters (which clustered in the middle of the UMAP) with low UMI and without expressing distinguishable markers for any cell type were removed. Some clusters such as neutrophils had lower numbers of genes expressed but showed distinct neutrophil markers. A cluster with a high number of doublets as indicated by the doublet scores was also removed. Differential gene expression between clusters was performed using the Wilcoxon test implemented in Seurat using FindAllMarkers or FindMarkers functions.

### Annotation and subsetting of the data

Small intestine data model pre-trained on the human ileal single data of the Human Gut Atlas was downloaded from the CellTypist website and was used to annotate the clusters with CellTypist.^65, 66^ Each of the 73 clusters were then classified into 8 different compartments - Mesenchymal, Myeloid, T/NK cells, B cells, plasma cells, epithelial cells, endothelial cells and EGCs based on the cell typist annotation. The annotation was additionally manually curated using canonical markers as shown in Figure 1D. Each compartment except EGCs were then separately re-clustered to reveal detailed heterogeneity. First re-clustering within each compartment at high resolution revealed low quality cells and doublets which were filtered out and only high-quality cells were retained for the second round of re-clustering.

Overall, an effort was made to stay consistent in annotation with earlier single cell atlas publication - the human gut atlas.^66^ For the mesenchymal cells and myeloid cells, Celltypist annotation based on human gut atlas was used wherever possible. For T cells, B cells, epithelial cells and plasma cells, manual annotation was performed after curating Celltypist annotations based on 2 publicly available annotated single cell RNA sequencing data sets of similar tissue. After re-clustering followed by filtering and annotation of subclusters in each compartment, all retained cells were used to compute the UMAP of all cells as in **Figure 1B**. The fine annotations of subclusters obtained from re-clustering each compartment was carried over to the metadata of all cells for downstream analyses such as CellPhoneDB.

### Gene regulatory network analysis

Gene regulatory network analysis was performed using the python implementation of single-cell regulatory network inference and clustering (SCENIC).^21^ Specifically, GRNBoost was used to construct a gene regulatory network from log normalised counts. Identified networks were then pruned using DNA motif analysis to remove indirect targets or associations and enrichment of each regulon in single cells were quantified using AUCell algorithm included in SCENIC. Further wilcoxon rank sum test as implemented in the Seurat R package was used to identify top significant transcription factors in each cluster. The analysis was performed using pySCENIC (0.11.1) - the python implementation of SCENIC.^67^

### Trajectory analysis

PAGA was used to estimate the connectivity between the Seurat clusters.^23^ 4 clusters including FAP fibroblasts identified by PAGA as connected were then used with Monocle3 to learn the differentiation trajectory.^25^ Healthy tissue specific GREM1^-^CD34^+^ fibroblasts were annotated as the root of the trajectory prior to ordering cells along the pseudotime. Similarly, for myeloid cell trajectory, PAGA was employed first to identify connected clusters. Connected clusters except pDCS were then used with monocle3 to construct the trajectory. Cells were ordered with CCR2 monocyte at the beginning of pseudotime.

### Intercellular interaction and signalling analysis

Overall ligand receptor interactions among cell compartments were analysed using CellPhoneDB.^27^ CellphoneDB analysis was separately performed for each of the 4 tissue segments on all cells using subcluster annotations. Specific ligands involved in altering gene expression leading to differentiation of FAP fibroblasts were done using the NicheNet R package with Kyoto Encyclopedia of Genes and Genomes (KEGG) database used for ligand-receptor pairs.^22, 28^ Genes differentially expressed between ABL2 fibroblasts, GREM1^-^CD34^+^ fibroblasts, GREM1^+^CD34^+^ fibroblasts, and FAP fibroblasts were considered as the geneset of interest. Visualisations were prepared using the ggplot2 R package except for Heatmap visualisations which were prepared using the heatmap R package.

### *In-silico* gene functional analysis

Functional analysis of up-regulated genes in specific clusters or a regulon was done using an enricher function from the ClusterProfiler package.^68, 69^ Also, for single cell data of myeloid and mesenchymal compartments, single sample gene set enrichment score (ssGSEA) as implemented in the scGSVA package was used with a combined database of KEGG, Reactome, Gene Ontology (GO), HALLMARK, BIOCARTA and Human Phenotype (HP).^22, 26, 70–72^ The Wilcoxon test was used to estimate the statistical significance of the term enrichment in the clusters. For specific custom geneset of Core matrisome collagens or core ECM genes, gene sets were downloaded from the matrisomeDB database, and the AddModuleScore function from Seurat package was used to create a module score at the single cell level.^73^

### Comparison of IBD data sets

For Seurat label transfer, we used our integrated mesenchymal data and myeloid data as references for the corresponding subsets of the 3 published IBD data sets.^14–16, 61^ To transfer cell type labels, we employed the Seurat label transfer method. The “FindTransferAnchors” function was applied to identify anchors between the reference and each of the published datasets, utilizing the CCA dimensionality reduction method. Subsequently, the “TransferData” function was used to transfer the labels from the reference dataset to the published datasets based on these anchors. Cell type classification probability was visualized with heatmaps.

To create a shared gene signature for FAP fibroblasts and Inflammatory monocytes in Inflammatory Bowel Disease (IBD), individual datasets were divided into two groups: myeloid cells (monocytes, macrophages, and dendritic cells) and mesenchymal cells (fibroblasts, myofibroblasts, and pericytes). The data were then analysed to identify clusters of cells that were similar to specific cell types such as FAP fibroblasts or Inflammatory monocytes. These subsets were annotated as Inflammatory fibroblasts or Inflammatory monocytes in various studies. Fibroblast subsets similar to FAP fibroblast were annotated as *Inflammatory fibroblasts* in Smillie et al., *activated fibroblasts* in Martin et al. and *Inflammatory fibroblasts IL11 CHI3L1* in Kong et al. Myeloid subsets similar to inflammatory monocytes were as annotated as *Inflammatory monocytes* in Smillie et al., *Inflammatory macrophages* in Martin et al. and *Monocytes S100A8 S100A9* in Kong et al.

Next, significantly up-regulated genes (with an adjusted p-value <0.05) in the Inflammatory monocytes or Inflammatory fibroblasts were identified within the respective compartments of each dataset using the FindMarkers function from Seurat. These lists of up-regulated genes were combined to create a common gene signature for Inflammatory monocytes or FAP fibroblasts in IBD (Table S4 and S5).

Finally, a single Seurat object with raw counts was created for each cell compartment (myeloid or mesenchymal), retaining the original annotation and meta-data, and normalised. The gene signature was then scored for each cell compartment using the AddModuleScore function from Seurat.

### Flow cytometry, sorting and analysis

To validate cell proportion of scRNA-seq in fibroblast subsets, cell suspensions were labelled with CD45-PE (1:300, clone 2D1, BioLegend), CD326-PE (1:300, clone 9C4, Biolegend), CD31-PE (1:300, clone WM59, Biolegend), CD90-BV421 (1:400, clone 5E10, Biolegend), podoplanin (PDPN)-APC (1:300, clone NC-08, Biolegend), CD34-FITC (1:200, clone 4H11, eBioscience), and FAP-Alexa fluor 750 (1:300, clone 427819, R&D Systems). 7-AAD (1:100, 559925, BD Biosciences) was used to determine cell viability. The combination of CD45, CD31 and CD326 was used as lineage to eliminate immune cells, endothelial cells and epithelial cells.

CD45-FITC (1:400, clone HI30, Biolegend), CD3-APC/Cy7 (1:300, clone UCHT1, Biolegend), CD56-APC/Cy7 (1:300, clone HCD56, Biolegend), CD19-APC/Cy7 (1:300, clone HIB19, Biolegend), HLA-DR-APC (1:300, clone L243, Biolegend), CD150 (SLAMF-1)-BV421 (1:100, clone A12, BD Horizon), CD14-PE/Cy7 (1:400, clone 63D3, Biolegend), CCR2-PE (1:400, clone LS132.1D9, BD Pharmingen) and CD206-PE/CF594 (1:300, clone 19.2, BD Horizon) were used to identify different subsets of myeloid cells. The combination of CD3, CD19 and CD56 was used as lineage to eliminate T cells, B cells and NK cells. Flow cytometry and sorting were performed on the MA900 multi-application cell sorter (Sony) with 100 µm sorting chips.

Sorted-HLA-DR^+^ cells, CD14^dim^ cells, CD150^-^CCR2^+^ cells and CD150^+^CCR2^+^ cells (10,000 cells per well) were seeded in a 96-well clear round bottom plate (3788, Corning) with 100 µL of RPMI-1640 medium, supplied with 5% FBS (BWSTS181H, VWR), 1% HEPES (15630056, Gibco), 1% L-glutamine (A2916801, Gibco), 1% sodium pyruvate (11360070, Gibco), and 1% antibiotic/antimycotic solution (A5955, Sigma-Aldrich) for 16 hours. The cell supernatants were collected and used to stimulate fibroblasts.

### RNA isolation and gene expression

RNA was extracted utilising the micro plus Kit (Qiagen), following the manufacturer’s guidelines. cDNA was then synthesised the High-Capacity cDNA Reverse Transcription Kit (ThermoFisher Scientific), as per the provided instructions. The RT-PCR process was carried out using the LightCycler 480 SYBR Green I Master (Roche) on a Light Cycler 480 instrument (Roche). The 2^−ΔCT^ method was employed to quantify the results, and gene expression levels of interest were normalised relative to the reference gene *RPLP0*.

### Primary human ileal fibroblasts and CCD18-Co colonic human fibroblast

Single-cell suspensions were directly cultured in a T-25 flask (90026, TPP) with RPMI-1640 medium (31870074, Gibco), supplied with 10% FBS (BWSTS181H, VWR), 1% HEPES (15630056, Gibco), 1% L-glutamine (A2916801, Gibco), 1% sodium pyruvate (11360070, Gibco), and 1% antibiotic/antimycotic solution (A5955, Sigma-Aldrich). 0.25% Trypsin-EDTA (25200056, Gibco) was used to detach the cells. Primary human ileal fibroblasts were obtained, purified and used after two passages. CCD18-Co colonic fibroblasts (CRL-1459) were obtained from ATCC and cultured according to the ATCC culture guides. Primary ileal fibroblasts and CCD-18Co fibroblasts were cultured in a 96-well plate for cell proliferation assay, scratch wound assay, *in vitro*-extracellular matrix deposition and immunofluorescent staining to determine fibroblast phenotypes and therapeutic function of Harmine (5µM, 286044, Sigma-Aldrich). Different combinations of 5ng/mL human IL-1α (R&D Systems), 5ng/mL human IL-1β (R&D Systems), 5ng/mL human TGFβ1 (R&D Systems), 5ng/mL human TNFα (R&D Systems), 5ng/mL human IFNγ (R&D Systems) or 5ng/mL human OSM (R&D Systems) were used to stimulate fibroblasts along with 5% FBS RMIP-1640 medium.

### *In vitro* immunofluorescence staining

CCD18-Co colonic human fibroblast (CRL-1459, ATCC) or primary ileal fibroblasts (10,000 cells per well) were cultured in the matrix-gel-coated PhenoPlate™ 96-well microplates (PerkinElmer) with inflammatory cytokines or myeloid derived supernatants along with Harmine (5µM, 286044, Sigma-Aldrich) for 24 or 48 hours. Then the cells were fixed with ice-cold acetone (Chem-Lab) and labelled with sheep anti-human FAP antibody (1:40, AF3715, R&D Systems), mouse anti-human TWIST1 antibody (1:100, clone 2F8E7, Invitrogen), goat anti-human collagen III alpha 1 antibody (1:500, NBP1-26547, Novus Biologicals), ATTO 490LS-conjugated phalloidin (1:200, 14479, Sigma-Aldrich), Alexa Fluor 488-conjugated donkey anti-sheep IgG antibody (A-11015, Invitrogen), Cy3-conjugated donkey anti-goat IgG antibody (705-165-003, Jackson ImmunoResearch) and Cy5-conjugated donkey anti-mouse IgG antibody (715-175-151, Jackson ImmunoResearch). Immunofluorescent staining was imaged on the Operetta CLS high-content analysis system (PerkinElmer) and analysed on Harmony 4.5 software (PerkinElmer) at the VIB Leuven.

### *In vitro*-extracellular matrix (ECM) deposition

To generate fibroblast-derived ECM, we modified the protocol of Kaukonen et al.^74^ Briefly, 0.2% (wt/vol) gelatin (G6144, Sigma-Aldrich)-coated PhenoPlate™ 96-well microplates (PerkinElmer) were treated with 1% (vol/vol) glutaraldehyde (G5882, Sigma-Aldrich) to create cross-link, following by 1 M glycine (1.04201, Millipore) to aspirate the cross-linker. CCD18- Co colonic human fibroblast (CRL-1459, ATCC) or primary ileal fibroblasts (30,000 cells per well) were seeded until 85-90% confluence. Fibroblasts were treated with/without inflammatory cytokines or with/without myeloid derived supernatants, described above for 72 hours, following by 50 µg/mL L-ascorbic acid (A4544, Sigma-Aldrich) treatment to promote collagen cross-linking daily. Fibroblasts were re-stimulated to reboot fibrotic phenotype every seven days. This cycle was repeated for three weeks till ECM was well deposited. Harmine (5 µM, 286044, Sigma-Aldrich) were given and refreshed daily after first stimulation.

ECM was labelled with primary antibodies, including goat anti-human collagen III alpha 1 antibody (1:500, NBP1-26547, Novus Biologicals), rabbit anti-human collagen I antibody (1:500, NB600-408, Novus Biologicals), mouse anti-human collagen IV antibody (1:500, clone 1042, eBioscience), sheep anti-human FAP antibody (1:40, AF3715, R&D Systems), and sheep anti-human fibronectin antibody (1:500, AF1918, R&D Systems) in cell culture medium for 1 hour in the cell culture incubator. Then the cells were labelled with secondary antibodies, including Alexa Fluor 488-conjugated donkey anti-mouse IgG antibody (715-545-150, Jackson ImmunoResearch), Cy3-conjugated donkey anti-goat IgG antibody (705-165-003, Jackson ImmunoResearch), Cy3-conjugated donkey anti-rabbit IgG antibody (711-165-152, Jackson ImmunoResearch) and Alexa Fluor 647-conjugated donkey anti-sheep IgG antibody (713-605-147, Jackson ImmunoResearch) before 4 % paraformaldehyde fixation (1040051000, Merck). ECM deposition was imaged on the LSM780 confocal microscope (Zeiss) with multiple focal planes (Z-stacks) and on the Operetta CLS high-content analysis system (PerkinElmer) and analysed on Harmony 4.5 software (PerkinElmer).

### Human-induced pluripotent stem cells (iPSCs) maintenance

Human-induced pluripotent stem cells (iPSCs) (WiCell, DF19-9-7T) were cultured on Cultrex Extract (3434-001-02, Stem Cell Qualified Reduced Growth Factor Basement Membrane, R&D Systems)-coated tissue culture dishes and were maintained in the mTeSR™ Plus medium (100-0274, STEMCELL Technologies). The medium was changed every two days. When the cells reached about 80% confluence, they were passaged using TrypLE™ Express Enzyme (12604013, Gibco), and the 10 μM RHO/ROCK pathway inhibitor (129830-38-2, STEMCELL Technologies) was used for 24 hours. The next day, the cells were washed with fresh medium and maintained until the next passage.

### Intestinal organoids (IOs) differentiation

Human intestinal organoids (IOs) were differentiated from iPSCs using an adapted protocol already published. The iPSCs were induced into 3D IOs in a three-step protocol. Briefly, on day 0, confluent iPSCs were induced into definitive endoderm (DE) differentiation using RPMI-1640 (31870074, Gibco) supplemented with Activin A (100 ng/mL, 338-AC, R&D Systems). The next day, the medium was changed and fresh RPMI-1640 (31870074, Gibco) supplemented with Activin A (100 ng/mL, 338-AC, R&D Systems) and 0.2% HyClone-defined FBS (GE Healthcare Bio-Sciences) was added. On the 3rd and 4th days, the medium was changed and RMPI-1640 (31870074, Gibco) supplemented with Activin A (100 ng/mL, 338-AC, R&D Systems) and 2 % HyClone-defined FBS (GE Healthcare Bio-Sciences) was added. Further, the DE was induced to mid- and hindgut differentiation by changing the medium daily (RPMI-1640 supplemented with 15 mM HEPES, 2% HyClone-defined FBS, FGF-4 (500 ng/mL, 235-F4, R&D Systems), WNT-3a (500 ng/mL, 5036-WN, R&D Systems) until the 3D spheroids were formed. Spheroids were collected and embedded in a drop Cultrex RGF Basement Membrane Extract, Type 2 (3536-005-02, R&D Systems) and fed with IOs complete medium (Advanced DMEM F12 (1264-010, Gibco) supplemented with B27 supplement (17504-044, Thermo Fisher Scientific), GluMAX supplement (35050-038, Thermo Fisher Scientific), penicillin and streptomycin (500 U/mL), R-spondin 1 (500 ng/mL, 4645-RS, R&D Systems), Noggin (100 ng/mL, 6057-NG, R&D Systems), EGF (100 ng/mL, 236-EG,R&D Systems). The medium was changed twice a week. After 50 days the IOs were used for the experiments.

### Immunofluorescence staining of IOs

IOs were washed with PBS and fixed with 4 % paraformaldehyde (PFA) for 20 minutes at room temperature and then washed three times with PBS. The IOs were dehydrated in 15 % sucrose at 4°C overnight. The IOs were frozen in Tissue Freezing Medium (Leica Biosystems). The samples were cut and rehydrated in PBS for 10 minutes. Subsequently, the samples were blocked with 1 % bovine serum albumin (BSA) at room temperature for 45 minutes and incubated with primary anti-FAP (1:200, F11-24, eBioscience), anti-PDPN (1:100, NZ-1.3, eBioscience), and anti-E-Cadherin (1:300, 24E10, Cell Signaling Technology) antibodies at 4 °C overnight. The samples were washed three times with PBS and then incubated with secondary antibodies, including Alexa Fluor 555-conjugated goat anti-mouse IgG antibody (A32727, Invitrogen), Alexa Fluor 488-conjugated donkey anti-rat IgG antibody (A-21208, Invitrogen), and Alexa Fluor 647-conjugated goat anti-rabbit IgG antibody (A-21245, Invitrogen) for one hour at room temperature. The samples were washed with PBS three times and stained with DAPI for twenty minutes at room temperature. The samples were washed with deionized water and embedded in Mowiol. The samples were acquired with Zeiss LSM 780 confocal microscope with 10x magnification and analysed with Fiji ImageJ.

### Chronic DSS colitis

Animal studies (project P188/2019) were performed under supervision of the animal ethics committee at the KU Leuven. Male C57BL/6 mice (aged 10 weeks, weight 28–30 gr) used for this study were group-housed under controlled temperature (22°C) and photoperiod (12∶12-hour light-dark cycles) conditions and given unrestricted access to standard rodent diet and water (or DSS drinking solution). The mice were subjected to 3 cycles of 2% DSS (w/v) (40 kDa, DS001, TdB Labs) administration (7 days of DSS administration followed by 14 days of regular drinking water was defined as one cycle). The mice were monitored for signs of colitis each day (i.e., body weight loss, diarrhoea score and blood in the faeces). Hemoccult II® Fecal Occult Blood Test Kit (61200, Beckman Coulter) was used to detect blood in faeces. Harmine (10 mg/kg) was dissolved in DMSO and Tween-80 and injected intraperitoneally from day 0 to day 10 during each DSS cycle. Mice were euthanized using CO^2^. Colon, spleen and kidney were taken for the analysis.

### Picrosirius red staining of the mouse colons

Mouse colon was fixed in 4% formaldehyde, and biopsies were embedded in paraffin and cut into 5µm thick sections for histological analysis. In brief, the tissue slides were de-waxed by HistoChoice clearing agent (H2779-1L, Sigma-Aldrich) and rehydrated by the gradient decrease of ethanol solution and water. Then the tissue was stained in 0.1% of Direct Red 80 (365548-5G, Sigma-Aldrich) in picric acid solution (P6744-1GA, Sigma-Aldrich) for an hour, followed by two changes of acidified water and 100% ethanol. To quantify the relative histologic area of collagen on Masson’s trichrome stained slides, we took an average of ten images in each sample on Leica DM2500 M and quantified them with Image J.

### Statistical analysis

Data are shown as mean ± standard error of the mean (SEM) with individual data values produced by using the GraphPad Prism V.9.1.0 software (GraphPad Inc). Multiple groups were compared by one-way analysis of variance (ANOVA) with Tukey’s multiple comparison test.

### Data and code availability

Single-cell RNA-seq data will be publicly available as of the date of publication. Microscopy data reported in this paper will be shared by the lead contact upon request. Any additional information required to re-analyze the data reported in this paper is available from the lead contact upon request.

---

**Figure S1.**
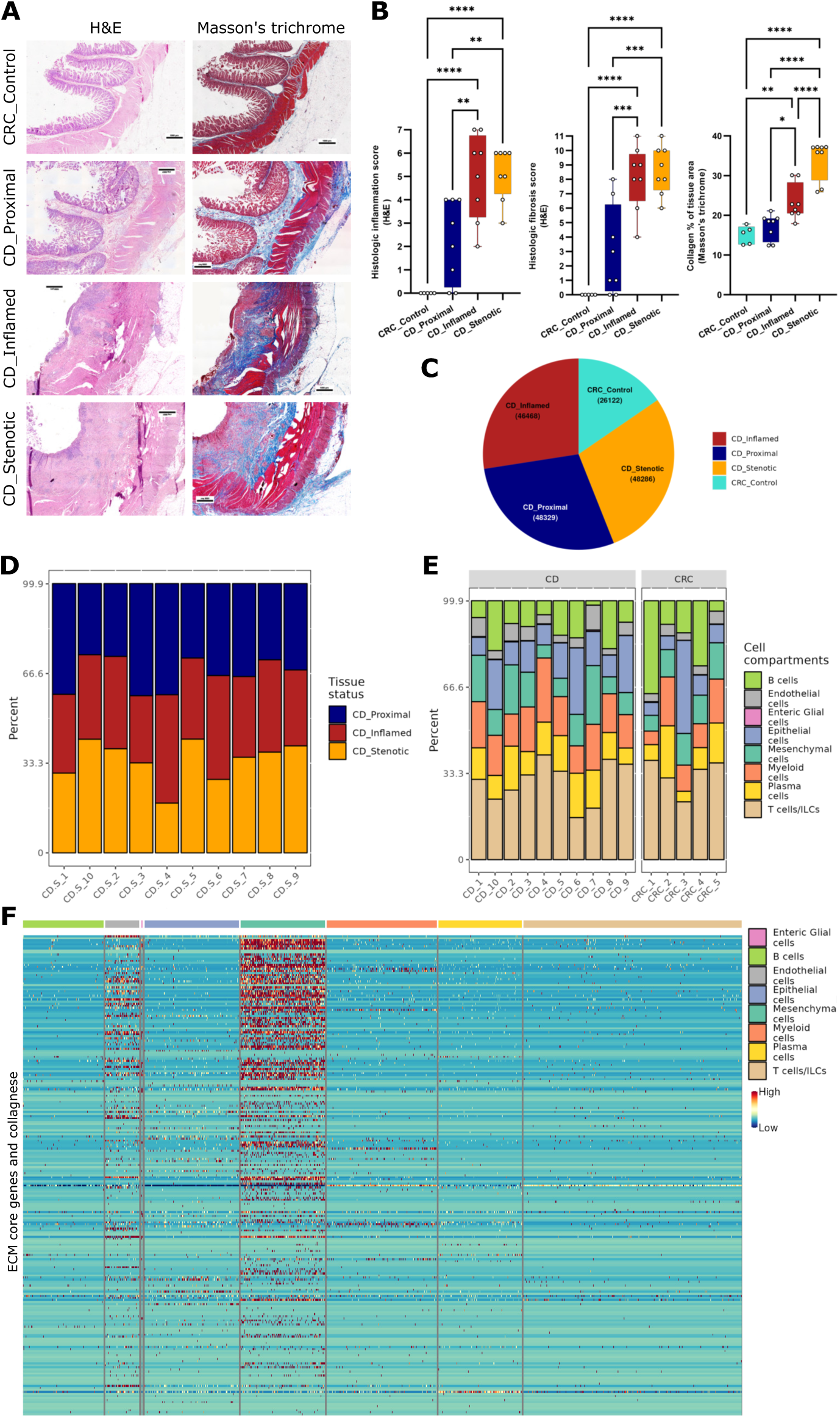
Single-cell profiling of fibro-stenotic ileal from CD and control ileum from CRC, related to Figure 1. (**A**) H&E and Masson’s trichrome stain showing signs of inflammation and fibrosis. (**B**) the plot of histologic scores and collagen quantification in different lesions of terminal ileum. Data are shown as box and whisker plots. Statistically significant differences were determined using a one-way ANOVA test corrected with Tukey’s multiple comparisons test (\**p* <0.05, ** *p* <0.01, *** *p* <0.005, \*\*\*\**p* <0.001). (**C**) Pie plot showing proportions of cells from different lesions of terminal ileum. (**D**) Bar plot showing cell proportions in different lesions in CD’s terminal ileum. (**E**) Bar plot showing percentage of cell clusters among the patients. (**F**) Heatmap showing core ECM genes in each cell type.

**Figure S2.**
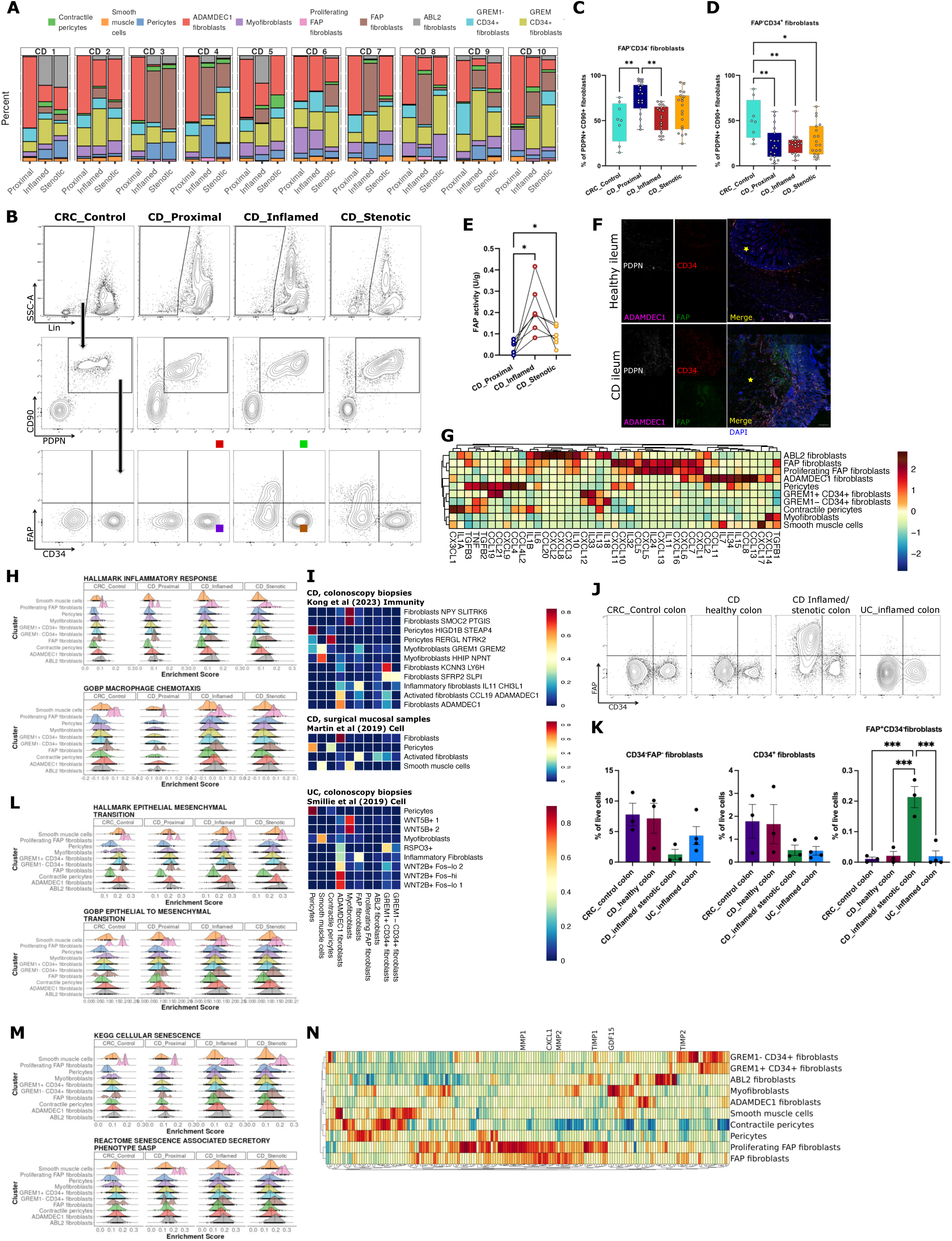
Heterogeneity of fibroblast in fibro-stenotic CD, related to Figure 2. (**A**) Bar plot showing proportions of mesenchymal subtypes in each terminal ileal lesion sample from each CD patient in the scRNA-seq data. (**B**) Contour plot showing gating strategy of fibroblast subsets by flow cytometry. (**C**) and (**D**) Percentage of fibroblast subsets in different lesions of terminal ileum are shown as bar plot with SEM. Statistically significant differences were determined using a one-way ANOVA test corrected with Tukey’s multiple comparisons test *(*p* <0.05*, **p* <0.01). (**E**) Before-after plot showing FAP activity in different lesions of terminal ileum. Data is shown as a before-after plot. Statistically significant differences were determined using a one-way ANOVA test corrected with Tukey’s multiple comparisons test (\**p* <0.05, \*\**p* <0.01). (**F**) Immunofluorescence staining for PDPN, ADAMDEC1, CD34 and FAP expression in healthy ileum and CD diseased ileum. Original image composed of stitched 25× images. (**G**) Heatmap depicting expression of chemokine and chemokine ligand genes in fibroblast subsets. (**H**) ssGSEA scores depicted as ridgeplots for selected terms, related to inflammation in fibroblast subsets. (**I**) Cross-dataset cell type prediction score heatmap showed similarity of stromal cell subset among published human IBD atlas. (**J**) Contour plot showing gating strategy of fibroblast subsets in colon by flow cytometry. (**K**) Percentage of fibroblast subsets in different lesions of colon are shown as bar plot with SEM. Statistically significant differences were determined using a one-way ANOVA test corrected with Tukey’s multiple comparisons test (\*\*\**p* <0.005). (**L**) ssGSEA scores depicted as ridgeplots for selected terms, related to EMT in fibroblast subsets. (**M**) ssGSEA scores depicted as ridgeplots for selected terms, related to cellular senescence in fibroblast subsets. (**N**) Heatmap showing relative expression of senescence-associated secretory phenotype (SASP) markers.

**Figure S3.**
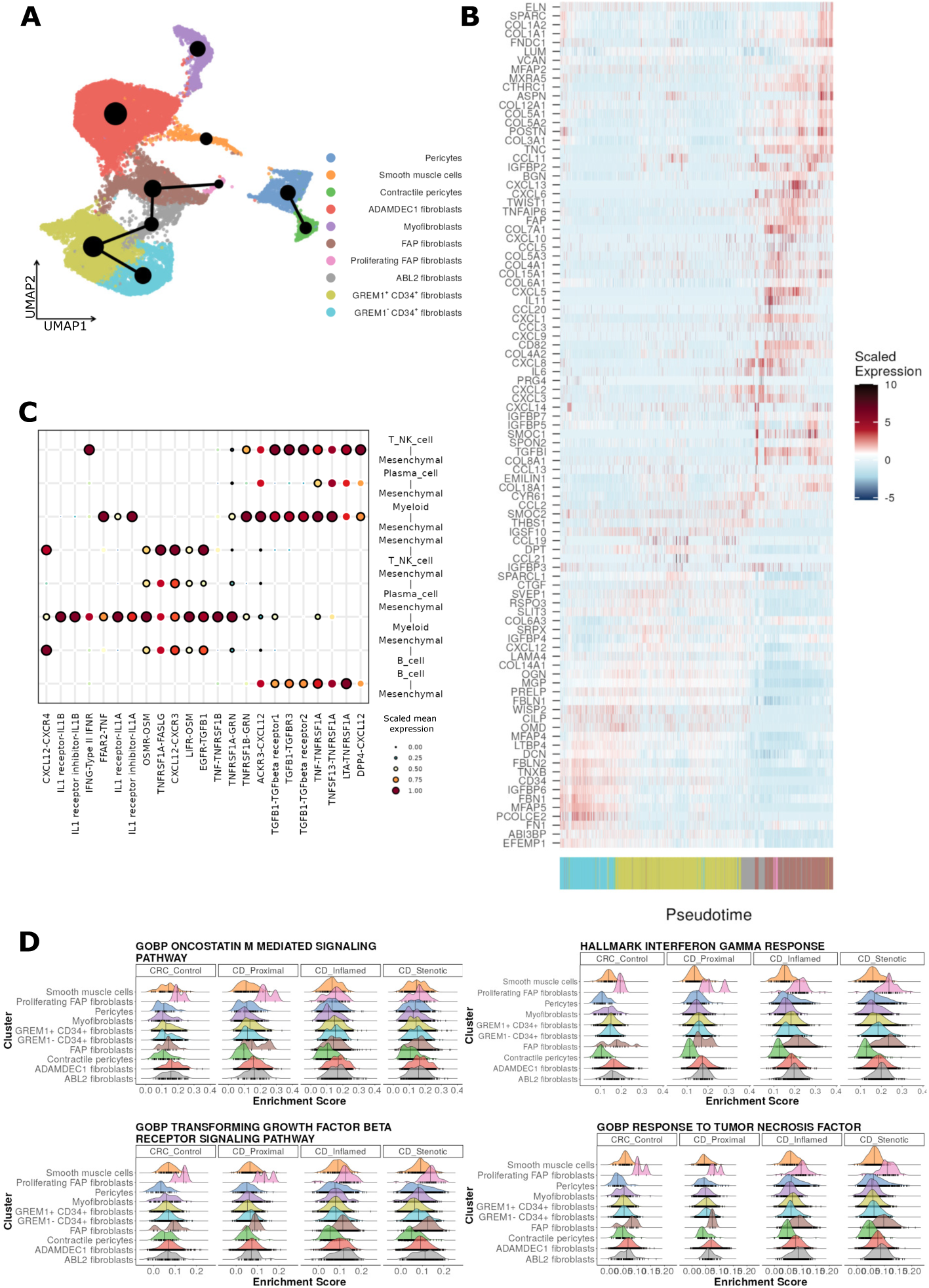
Fibroblast-myeloid cell interaction modulates fibro-stenosis, related to Figure 3. (**A**) Partition-based graph abstraction (PAGA) analysis on fibroblast subsets showing most likely trajectories. (**B**) Heatmap showing gene expression change along cell Monocle3 pseudo-time. (**C**) Cellphone DB dot plot showing ligand-receptor interactions between mesenchymal compartment and immune cells. First and second interacting molecules on x-axis correspond to first and second cell types on the y-axis respectively. Black circles indicate significant interactions (**D**) ssGSEA scores depicted as ridgeplots for selected terms in fibroblast subsets.

**Figure S4.**
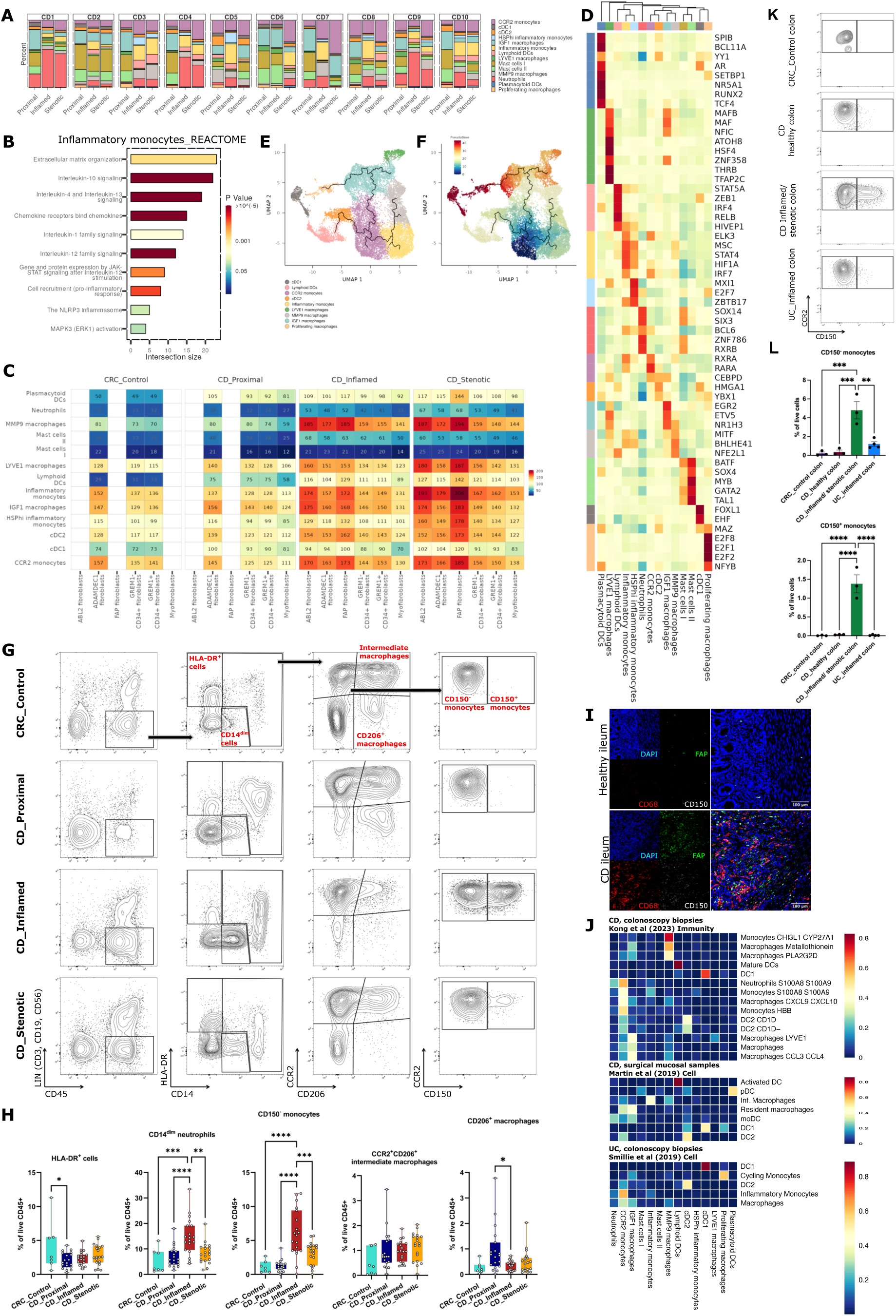
Heterogeneity of myeloid cells in fibro-stenotic CD, related to Figure 4. **(A)** Bar plot showing proportion of myeloid subsets in each terminal ileal lesion sample from each CD patient in the scRNA-seq data. (**B**) Selected terms from the Reactome biological pathway enrichment analysis for differentially upregulated genes in Inflammatory monocytes (logFC,>0.5; FDR, <0.1). (**C**) Heatmap showing number of interactions (Ligand-Receptor pairs) between myeloid and mesenchymal. (**D**) SCENIC showing relative transcription factor activity in each myeloid cell subset. (**E**) and (**F**) Pseudo-time trajectory projected onto a UMAP of selected myeloid subsets. (**G**) Contour plot showing gating strategy of myeloid subsets by flow cytometry among the different disease states. (**H**) Bar plots showing cell proportion of myeloid subsets in different lesions of terminal ileum. Data are shown as bar plots with SEM. Statistically significant differences were determined using a one-way ANOVA test corrected with Tukey’s multiple comparisons test (\**p* <0.05, ** *p* <0.01, *** *p* <0.005, **** *p* <0.001). (**I**) Immunofluorescence staining for CD68, SLAMF1 (CD150) and FAP expression in healthy ileum and CD diseased ileum. Original image composed of stitched 25× images. (**J**) Cross-dataset cell type prediction score heatmap showed similarity of myeloid cell subset among published human IBD atlas. (**K**) Contour plot showing gating strategy of myeloid subsets by flow cytometry among the colon from CD and UC patients. (**L**) Bar plots showing cell proportion of CCR2 monocytes and CD150 Inflammatory monocytes in different lesions of colon. Data are shown as bar plots with SEM. Statistically significant differences were determined using a one-way ANOVA test corrected with Tukey’s multiple comparisons test (\*\**p* <0.01, *** *p* <0.005, **** *p* <0.001).

**Figure S5.**
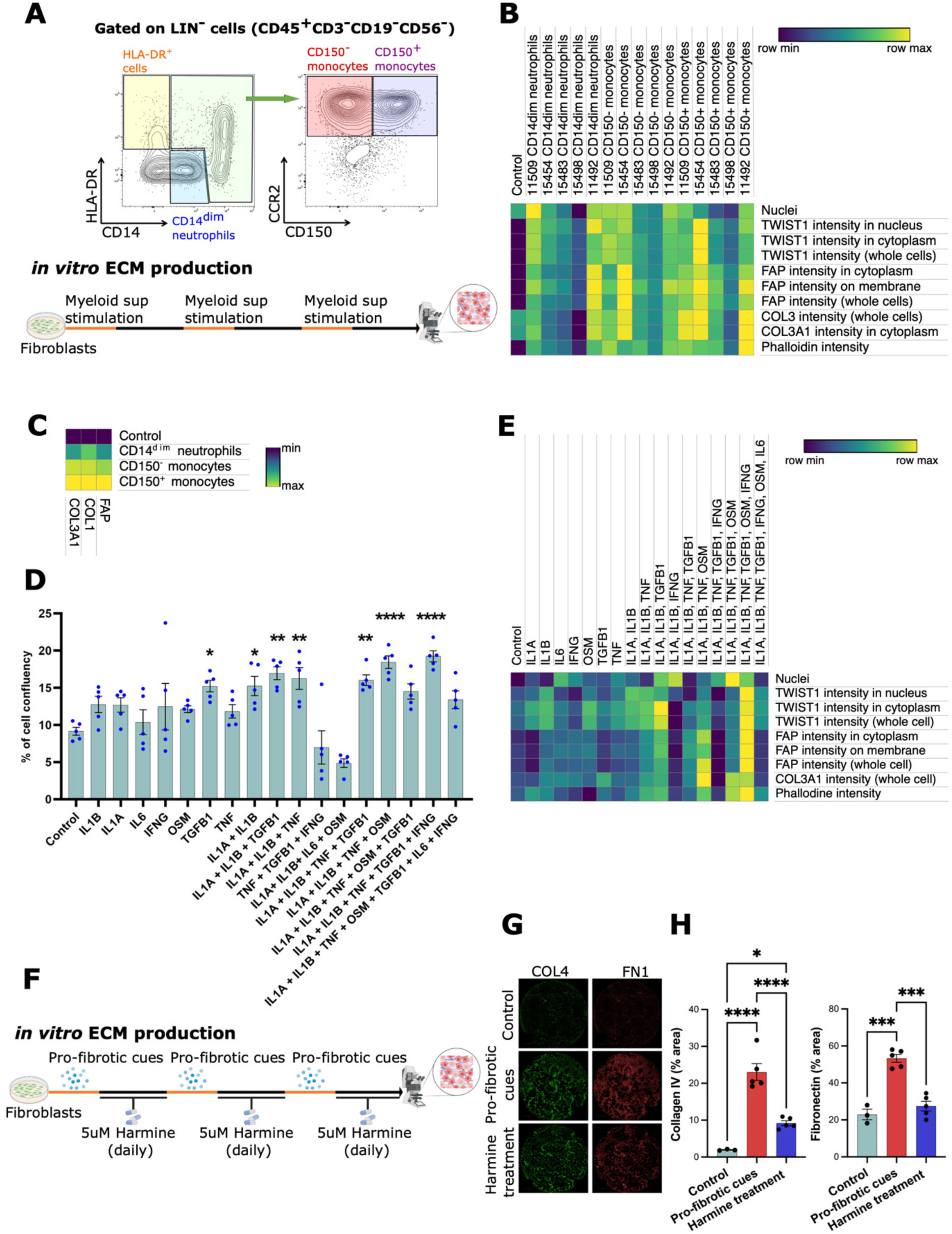
CD150^+^ monocytes-derived cytokines promoted FAP fibroblast activation and extra-cellular matrix protein deposition, related to Figure 5. (**A**) Experimental workflow showing FACS-sorting gating strategy and setup for *in vitro* ECM production. (**B**) Heatmap showing relative expression of FAP, TWIST1 and type III collagen in CCD-18Co fibroblast treated with supernatant from FACS sorted myeloid subsets (n= 5 individual patients). (**C**) Heatmap showing extracellular deposition of FAP, type I collagen and type III collagen in monocyte-stimulated CCD-18Co fibroblasts. (**D**) Heatmap showing relative protein expression of FAP, TWIST1 and type II collagen in primary ileal fibroblasts after being stimulated by selected cytokine combinations, predicted by NicheNet. (**E**) Experimental workflow. (**F**) Immunofluorescence staining (10× image) and (**G**) Bar plot showing relative expression of fibronectin and type IV collagen in *pro-fibrotic cues*-stimulated CCD-18Co fibroblasts after TWIST1 inhibition. Data are shown as bar plots with SEM. Statistically significant differences were determined using a one-way ANOVA test corrected with Tukey’s multiple comparisons test (\**p* <0.05, ** *p* <0.01, *** *p* <0.005, **** *p* <0.001).

**Figure S6.**
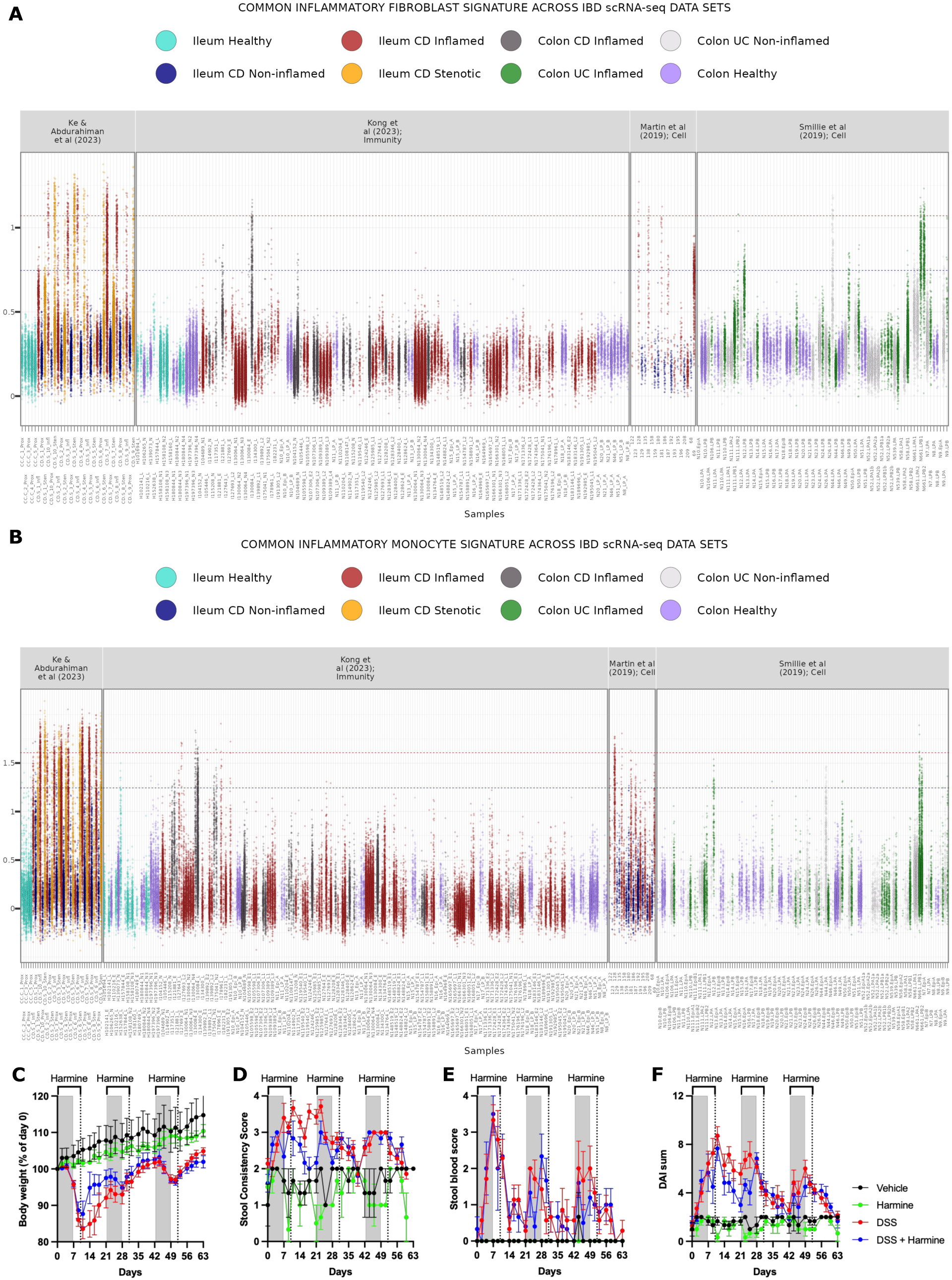
Previously published datasets using mucosal biopsies do not show consistent presence of FAP fibroblasts and inflammatory monocytes across IBD patients and TWIST1 inhibition attenuated intestinal fibrosis, related to Figure S2, S4 and 6. (**A**) Dotplot showing gene signature score for common markers of inflammatory fibroblasts in 4 independent IBD data sets. The area between the two horizontal lines indicates the interquantile range of the gene signature score for the cells in Ke & Abdurahiman et al. (**B**) Dotplot showing gene signature score for common markers of inflammatory monocytes in 4 independent IBD data sets. The area between the two horizontal lines indicates the interquantile range of the gene signature scores for the cells in Ke & Abdurahiman et al. (**C**) Body weight, (**D**) stool consistency, (**E**) stool blood and (**F**) disease activity index changing during 3 cycles of DSS colitis. Data are shown as X-Y plots with SEM. Statistically significant differences were determined using a one-way ANOVA test corrected with Tukey’s multiple comparisons test (\**p* <0.05, ** *p* <0.01, *** *p* <0.005, **** *p* <0.001).

## REFERENCES

1. Ng, S.C., Shi, H.Y., Hamidi, N., Underwood, F.E., Tang, W., Benchimol, E.I., Panaccione, R., Ghosh, S., Wu, J.C.Y., Chan, F.K.L., et al. (2017). Worldwide incidence and prevalence of inflammatory bowel disease in the 21st century: a systematic review of population-based studies. The Lancet 390, 2769–2778. 10.1016/S0140-6736(17)32448-0.

2. Kaplan, G.G. (2015). The global burden of IBD: from 2015 to 2025. Nat Rev Gastroenterol Hepatol 12, 720–727. 10.1038/nrgastro.2015.150.

3. Kappelman, M.D., Porter, C.Q., Galanko, J.A., Rifas-Shiman, S.L., Ollendorf, D.A., Sandler, R.S., and Finkelstein, J.A. (2011). Utilization of healthcare resources by U.S. children and adults with inflammatory bowel disease. Inflamm Bowel Dis 17, 62–68. 10.1002/ibd.21371.

4. Carter, M.J. (2004). Guidelines for the management of inflammatory bowel disease in adults. Gut 53, v1–v16. 10.1136/gut.2004.043372.

5. Frolkis, A.D., Dykeman, J., Negrón, M.E., Debruyn, J., Jette, N., Fiest, K.M., Frolkis, T., Barkema, H.W., Rioux, K.P., Panaccione, R., et al. (2013). Risk of surgery for inflammatory bowel diseases has decreased over time: A systematic review and meta-analysis of population-based studies. Gastroenterology 145, 996–1006. 10.1053/j.gastro.2013.07.041.

6. Lee, H.S., Park, S.K., and Park, D. Il (2018). Novel treatments for inflammatory bowel disease. Korean J Intern Med 33, 20. 10.3904/KJIM.2017.393.

7. Present, D.H., Rutgeerts, P., Targan, S., Hanauer, S.B., Mayer, L., van Hogezand, R.A., Podolsky, D.K., Sands, B.E., Braakman, T., DeWoody, K.L., et al. (1999). Infliximab for the Treatment of Fistulas in Patients with Crohn’s Disease. New England Journal of Medicine. 10.1056/nejm199905063401804.

8. Neurath, M.F. (2017). Current and emerging therapeutic targets for IBD. Nature Reviews Gastroenterology & Hepatology 2017 14:5 14, 269–278. 10.1038/nrgastro.2016.208.

9. Raine, T., and Danese, S. (2022). Breaking Through the Therapeutic Ceiling: What Will It Take? Gastroenterology 162, 1507–1511. 10.1053/J.GASTRO.2021.09.078.

10. Murthy, S.K., Begum, J., Benchimol, E.I., Bernstein, C.N., Kaplan, G.G., McCurdy, J.D., Singh, H., Targownik, L., and Taljaard, M. (2020). Introduction of anti-TNF therapy has not yielded expected declines in hospitalisation and intestinal resection rates in inflammatory bowel diseases: a population-based interrupted time series study. Gut 69, 274–282. 10.1136/GUTJNL-2019-318440.

11. Granlund, A. van B., Flatberg, A., Østvik, A.E., Drozdov, I., Gustafsson, B., Kidd, M., Beisvag, V., Torp, S.H., Waldum, H.L., Martinsen, T.C., et al. (2013). Whole Genome Gene Expression Meta-Analysis of Inflammatory Bowel Disease Colon Mucosa Demonstrates Lack of Major Differences between Crohn’s Disease and Ulcerative Colitis. PLoS One 8, e56818. 10.1371/JOURNAL.PONE.0056818.

12. Pelia, R., Venkateswaran, S., Matthews, J.D., Haberman, Y., Cutler, D.J., Hyams, J.S., Denson, L.A., and Kugathasan, S. (2021). Profiling non-coding RNA levels with clinical classifiers in pediatric Crohn’s disease. BMC Med Genomics 14, 194. 10.1186/S12920-021-01041-7.

13. Haberman, Y., Minar, P., Karns, R., Dexheimer, P.J., Ghandikota, S., Tegge, S., Shapiro, D., Shuler, B., Venkateswaran, S., Braun, T., et al. (2021). Mucosal Inflammatory and Wound Healing Gene Programmes Reveal Targets for Stricturing Behaviour in Paediatric Crohn’s Disease. J Crohns Colitis 15, 273–286. 10.1093/ECCO-JCC/JJAA166.

14. Martin, J.C., Chang, C., Boschetti, G., Ungaro, R., Giri, M., Grout, J.A., Gettler, K., Chuang, L. shiang, Nayar, S., Greenstein, A.J., et al. (2019). Single-Cell Analysis of Crohn’s Disease Lesions Identifies a Pathogenic Cellular Module Associated with Resistance to Anti-TNF Therapy. Cell 178, 1493–1508.e20. 10.1016/j.cell.2019.08.008.

15. Kong, L., Pokatayev, V., Lefkovith, A., Carter, G.T., Creasey, E.A., Krishna, C., Subramanian, S., Kochar, B., Ashenberg, O., Lau, H., et al. (2023). The landscape of immune dysregulation in Crohn’s disease revealed through single-cell transcriptomic profiling in the ileum and colon. Immunity 56, 444–458.e5. 10.1016/j.immuni.2023.01.002.

16. Smillie, C.S., Biton, M., Ordovas-Montanes, J., Sullivan, K.M., Burgin, G., Graham, D.B., Herbst, R.H., Rogel, N., Slyper, M., Waldman, J., et al. (2019). Intra- and Inter-cellular Rewiring of the Human Colon during Ulcerative Colitis. Cell 178, 714–730.e22. 10.1016/J.CELL.2019.06.029.

17. Gordon, I.O., Bettenworth, D., Bokemeyer, A., Srivastava, A., Rosty, C., de Hertogh, G., Robert, M.E., Valasek, M.A., Mao, R., Kurada, S., et al. (2020). Histopathology Scoring Systems of Stenosis Associated With Small Bowel Crohn’s Disease: A Systematic Review. Gastroenterology 158, 137–150.e1. 10.1053/J.GASTRO.2019.08.033.

18. Chen, W., Lu, C., Hirota, C., Iacucci, M., Ghosh, S., and Gui, X. (2017). Smooth Muscle Hyperplasia/Hypertrophy is the Most Prominent Histological Change in Crohn’s Fibrostenosing Bowel Strictures: A Semiquantitative Analysis by Using a Novel Histological Grading Scheme. J Crohns Colitis 11, 92–104. 10.1093/ECCO-JCC/JJW126.

19. Jansen, K., Heirbaut, L., Verkerk, R., Cheng, J.D., Joossens, J., Cos, P., Maes, L., Lambeir, A.M., De Meester, I., Augustyns, K., et al. (2014). Extended structure-activity relationship and pharmacokinetic investigation of (4-quinolinoyl)glycyl-2-cyanopyrrolidine inhibitors of fibroblast activation protein (FAP). J Med Chem 57, 3053–3074. 10.1021/JM500031W/SUPPL_FILE/JM500031W_SI_001.PDF.

20. Foroutan, M., Bhuva, D.D., Lyu, R., Horan, K., Cursons, J., and Davis, M.J. (2018). Single sample scoring of molecular phenotypes. BMC Bioinformatics 19. 10.1186/S12859-018-2435-4.

21. Aibar, S., González-Blas, C.B., Moerman, T., Huynh-Thu, V.A., Imrichova, H., Hulselmans, G., Rambow, F., Marine, J.-C., Geurts, P., Aerts, J., et al. (2017). SCENIC: single-cell regulatory network inference and clustering. Nat Methods 14, 1083–1086. 10.1038/nmeth.4463.

22. Kanehisa, M., Sato, Y., Kawashima, M., Furumichi, M., and Tanabe, M. (2016). KEGG as a reference resource for gene and protein annotation. Nucleic Acids Res 44, D457–D462. 10.1093/NAR/GKV1070.

23. Wolf, F.A., Hamey, F.K., Plass, M., Solana, J., Dahlin, J.S., Göttgens, B., Rajewsky, N., Simon, L., and Theis, F.J. (2019). PAGA: graph abstraction reconciles clustering with trajectory inference through a topology preserving map of single cells. Genome Biol 20, 1–9. 10.1186/S13059-019-1663-X/FIGURES/4.

24. Saul, D., Kosinsky, R.L., Atkinson, E.J., Doolittle, M.L., Zhang, X., LeBrasseur, N.K., Pignolo, R.J., Robbins, P.D., Niedernhofer, L.J., Ikeno, Y., et al. (2022). A new gene set identifies senescent cells and predicts senescence-associated pathways across tissues. Nat Commun 13. 10.1038/S41467-022-32552-1.

25. Cao, J., Spielmann, M., Qiu, X., Huang, X., Ibrahim, D.M., Hill, A.J., Zhang, F., Mundlos, S., Christiansen, L., Steemers, F.J., et al. (2019). The single cell transcriptional landscape of mammalian organogenesis. Nature 566, 496. 10.1038/S41586-019-0969-X.

26. Gillespie, M., Jassal, B., Stephan, R., Milacic, M., Rothfels, K., Senff-Ribeiro, A., Griss, J., Sevilla, C., Matthews, L., Gong, C., et al. (2022). The reactome pathway knowledgebase 2022. Nucleic Acids Res 50, D687–D692. 10.1093/NAR/GKAB1028.

27. Efremova, M., Vento-Tormo, M., Teichmann, S.A., and Vento-Tormo, R. (2020). CellPhoneDB: inferring cell-cell communication from combined expression of multi-subunit ligand-receptor complexes. Nat Protoc 15, 1484–1506. 10.1038/S41596-020-0292-X.

28. Browaeys, R., Saelens, W., and Saeys, Y. (2019). NicheNet: modeling intercellular communication by linking ligands to target genes. Nature Methods 2019 17:2 *17*, 159–162. 10.1038/s41592-019-0667-5.

29. Smedh, K., Olaison, G., Franzén, L., and Sjödahl, R. (1996). The endoscopic picture reflects transmural inflammation better than endoscopic biopsy in Crohn’s disease. Eur J Gastroenterol Hepatol 8, 1189–1193. 10.1097/00042737-199612000-00011.

30. Stzepourginski, I., Nigro, G., Jacob, J.-M., Dulauroy, S., Sansonetti, P.J., Eberl, G., and Peduto, L. (2017). CD34+ mesenchymal cells are a major component of the intestinal stem cells niche at homeostasis and after injury. Proceedings of the National Academy of Sciences 114, E506– E513. 10.1073/PNAS.1620059114.

31. Mulder, K., Patel, A.A., Kong, W.T., Piot, C., Halitzki, E., Dunsmore, G., Khalilnezhad, S., Irac, S.E., Dubuisson, A., Chevrier, M., et al. (2021). Cross-tissue single-cell landscape of human monocytes and macrophages in health and disease. Immunity 54, 1883–1900.e5. 10.1016/J.IMMUNI.2021.07.007.

32. Du, L., Sun, X., Gong, H., Wang, T., Jiang, L., Huang, C., Xu, X., Li, Z., Xu, H., Ma, L., et al. (2023). Single cell and lineage tracing studies reveal the impact of CD34+ cells on myocardial fibrosis during heart failure. Stem Cell Res Ther 14, 1–25. 10.1186/S13287-023-03256-0/FIGURES/8.

33. Ansieau, S., Morel, A.P., Hinkal, G., Bastid, J., and Puisieux, A. (2010). TWISTing an embryonic transcription factor into an oncoprotein. Oncogene 29, 3173–3184. 10.1038/ONC.2010.92.

34. Yeung, K.T., and Yang, J. (2017). Epithelial–mesenchymal transition in tumor metastasis. Mol Oncol 11, 28. 10.1002/1878-0261.12017.

35. Ning, X., Zhang, K., Wu, Q., Liu, M., and Sun, S. (2018). Emerging role of Twist1 in fibrotic diseases. J Cell Mol Med 22, 1383. 10.1111/JCMM.13465.

36. García-Palmero, I., Torres, S., Bartolomé, R.A., Peláez-García, A., Larriba, M.J., Lopez-Lucendo, M., Peña, C., Escudero-Paniagua, B., Muñoz, A., and Casal, J.I. (2016). Twist1-induced activation of human fibroblasts promotes matrix stiffness by upregulating palladin and collagen α1(VI). Oncogene 35, 5224–5236. 10.1038/ONC.2016.57.

37. Liu, C., Mo, L.H., Feng, B.S., Jin, Q.R., Li, Y., Lin, J., Shu, Q., Liu, Z.G., Liu, Z., Sun, X., et al. (2021). Twist1 contributes to developing and sustaining corticosteroid resistance in ulcerative colitis. Theranostics 11, 7797–7812. 10.7150/THNO.62256.

38. Guilliams, M., Thierry, G.R., Bonnardel, J., and Bajenoff, M. (2020). Establishment and Maintenance of the Macrophage Niche. Immunity 52, 434–451. 10.1016/J.IMMUNI.2020.02.015.

39. Nafie, E., Lolarga, J., Lam, B., Guo, J., Abdollahzadeh, E., Rodriguez, S., Glackin, C., and Liu, J. (2021). Harmine inhibits breast cancer cell migration and invasion by inducing the degradation of Twist1. PLoS One 16, e0247652. 10.1371/JOURNAL.PONE.0247652.

40. Wynn, T.A., and Vannella, K.M. (2016). Macrophages in Tissue Repair, Regeneration, and Fibrosis. Immunity 44, 450–462. 10.1016/J.IMMUNI.2016.02.015.

41. Yochum, Z.A., Cades, J., Mazzacurati, L., Neumann, N.M., Khetarpal, S.K., Chatterjee, S., Wang, H., Attar, M.A., Huang, E.H.B., Chatley, S.N., et al. (2017). A First-in-Class TWIST1 Inhibitor with Activity in Oncogene-Driven Lung Cancer. Mol Cancer Res 15, 1764–1776. 10.1158/1541-7786.MCR-17-0298.

42. Frucht, D.M., Aringer, M., Galon, J., Danning, C., Brown, M., Fan, S., Centola, M., Wu, C.-Y., Yamada, N., El Gabalawy, H., et al. (2000). Stat4 Is Expressed in Activated Peripheral Blood Monocytes, Dendritic Cells, and Macrophages at Sites of Th1-Mediated Inflammation. The Journal of Immunology 164, 4659–4664. 10.4049/JIMMUNOL.164.9.4659.

43. Foell, D., Wittkowski, H., Kessel, C., Lüken, A., Weinhage, T., Varga, G., Vogl, T., Wirth, T., Viemann, D., Björk, P., et al. (2013). Proinflammatory S100A12 can activate human monocytes via toll-like receptor 4. Am J Respir Crit Care Med 187, 1324–1334. 10.1164/RCCM.201209-1602OC/SUPPL_FILE/DISCLOSURES.PDF.

44. Schenk, M., Bouchon, A., Seibold, F., and Mueller, C. (2007). TREM-1--expressing intestinal macrophages crucially amplify chronic inflammation in experimental colitis and inflammatory bowel diseases. J Clin Invest 117, 3097–3106. 10.1172/JCI30602.

45. Frede, S., Stockmann, C., Freitag, P., and Fandrey, J. (2006). Bacterial lipopolysaccharide induces HIF-1 activation in human monocytes via p44/42 MAPK and NF-κB. Biochemical Journal 396, 517–527. 10.1042/BJ20051839.

46. Gaiani, F., Rotoli, B.M., Ferrari, F., Barilli, A., Visigalli, R., Carra, M.C., de’Angelis, G.L., de’Angelis, N., and Dall’Asta, V. (2020). Monocytes from infliximab-resistant patients with Crohn’s disease exhibit a disordered cytokine profile. Scientific Reports 2020 10:1 10, 1–10. 10.1038/s41598-020-68993-1.

47. Yang, A.-T., Kim, Y.-O., Yan, X.-Z., Abe, H., Aslam, M., Park, K.-S., Zhao, X.-Y., Jia, J.-D., Klein, T., You, H., et al. (2023). Fibroblast Activation Protein Activates Macrophages and Promotes Parenchymal Liver Inflammation and Fibrosis. Cell Mol Gastroenterol Hepatol 15, 841–867. 10.1016/J.JCMGH.2022.12.005.

48. Rurik, J.G., Tombácz, I., Yadegari, A., Méndez Fernández, P.O., Shewale, S. V., Li, L., Kimura, T., Soliman, O.Y., Papp, T.E., Tam, Y.K., et al. (2022). CAR T cells produced in vivo to treat cardiac injury. Science (1979) 375, 91–96. 10.1126/SCIENCE.ABM0594/SUPPL_FILE/SCIENCE.ABM0594_MDAR_REPRODUCIBILITY_CHECKLIST.PDF.

49. Tillmanns, J., Hoffmann, D., Habbaba, Y., Schmitto, J.D., Sedding, D., Fraccarollo, D., Galuppo, P., and Bauersachs, J. (2015). Fibroblast activation protein alpha expression identifies activated fibroblasts after myocardial infarction. J Mol Cell Cardiol 87, 194–203. 10.1016/j.yjmcc.2015.08.016.

50. Aghajanian, H., Kimura, T., Rurik, J.G., Hancock, A.S., Leibowitz, M.S., Li, L., Scholler, J., Monslow, J., Lo, A., Han, W., et al. (2019). Targeting cardiac fibrosis with engineered T cells. Nature 2019 573:7774 *573*, 430–433. 10.1038/s41586-019-1546-z.

51. Croft, A.P., Campos, J., Jansen, K., Turner, J.D., Marshall, J., Attar, M., Savary, L., Wehmeyer, C., Naylor, A.J., Kemble, S., et al. (2019). Distinct fibroblast subsets drive inflammation and damage in arthritis. Nature 2019 570:7760 570, 246–251. 10.1038/s41586-019-1263-7.

52. Kimura, T., Monslow, J., Klampatsa, A., Leibowitz, M., Sun, J., Liousia, M., Woodruff, P., Moon, E., Todd, L., Puré, E., et al. (2019). Loss of cells expressing fibroblast activation protein has variable effects in models of TGF-β and chronic bleomycin-induced fibrosis. Am J Physiol Lung Cell Mol Physiol 317, L271–L282. 10.1152/AJPLUNG.00071.2019/ASSET/IMAGES/LARGE/ZH50081976670006.JPEG.

53. Qi, J., Sun, H., Zhang, Y., Wang, Z., Xun, Z., Li, Z., Ding, X., Bao, R., Hong, L., Jia, W., et al. (2022). Single-cell and spatial analysis reveal interaction of FAP+ fibroblasts and SPP1+ macrophages in colorectal cancer. Nature Communications 2022 13:1 13, 1–20. 10.1038/s41467-022-29366-6.

54. Feig, C., Jones, J.O., Kraman, M., Wells, R.J.B., Deonarine, A., Chan, D.S., Connell, C.M., Roberts, E.W., Zhao, Q., Caballero, O.L., et al. (2013). Targeting CXCL12 from FAP-expressing carcinoma-associated fibroblasts synergizes with anti-PD-L1 immunotherapy in pancreatic cancer. Proc Natl Acad Sci U S A 110, 20212–20217. 10.1073/PNAS.1320318110/SUPPL_FILE/SAPP.PDF.

55. Habermann, A.C., Gutierrez, A.J., Bui, L.T., Yahn, S.L., Winters, N.I., Calvi, C.L., Peter, L., Chung, M.I., Taylor, C.J., Jetter, C., et al. (2020). Single-cell RNA sequencing reveals profibrotic roles of distinct epithelial and mesenchymal lineages in pulmonary fibrosis. Sci Adv 6. 10.1126/SCIADV.ABA1972/SUPPL_FILE/ABA1972_TABLE_S9.XLSX.

56. Friedrich, M., Pohin, M., Jackson, M.A., Korsunsky, I., Bullers, S.J., Rue-Albrecht, K., Christoforidou, Z., Sathananthan, D., Thomas, T., Ravindran, R., et al. (2021). IL-1-driven stromal–neutrophil interactions define a subset of patients with inflammatory bowel disease that does not respond to therapies. Nature Medicine 2021 27:11 27, 1970–1981. 10.1038/s41591-021-01520-5.

57. Bracke, A., Van Elzen, R., Van Der Veken, P., Augustyns, K., De Meester, I., and Lambeir, A.M. (2019). The development and validation of a combined kinetic fluorometric activity assay for fibroblast activation protein alpha and prolyl oligopeptidase in plasma. Clinica Chimica Acta 495, 154–160. 10.1016/J.CCA.2019.04.063.

58. McGinnis, C.S., Murrow, L.M., and Gartner, Z.J. (2019). DoubletFinder: Doublet Detection in Single-Cell RNA Sequencing Data Using Artificial Nearest Neighbors. Cell Syst 8, 329–337.e4. 10.1016/J.CELS.2019.03.003.

59. Ahlmann-Eltze, C., and Huber, W. (2021). glmGamPoi: fitting Gamma-Poisson generalized linear models on single cell count data. Bioinformatics 36, 5701–5702. 10.1093/BIOINFORMATICS/BTAA1009.

60. Wolock, S.L., Lopez, R., and Klein, A.M. (2019). Scrublet: Computational Identification of Cell Doublets in Single-Cell Transcriptomic Data. Cell Syst 8, 281–291.e9. 10.1016/J.CELS.2018.11.005.

61. Stuart, T., Butler, A., Hoffman, P., Hafemeister, C., Papalexi, E., Mauck, W.M., Hao, Y., Stoeckius, M., Smibert, P., and Satija, R. (2019). Comprehensive Integration of Single-Cell Data. Cell 177, 1888–1902.e21.10.1016/J.CELL.2019.05.031/ATTACHMENT/2F8B9EBE-54E6-43EB-9EF2-949B6BDA8BA2/MMC3.PDF.

62. Germain, P.L., Robinson, M.D., Lun, A., Garcia Meixide, C., and Macnair, W. (2021). Doublet identification in single-cell sequencing data using scDblFinder. F1000Res 10. 10.12688/F1000RESEARCH.73600.2.

63. Hafemeister, C., and Satija, R. (2019). Normalization and variance stabilization of single-cell RNA-seq data using regularized negative binomial regression. Genome Biol 20, 1–15. 10.1186/S13059-019-1874-1/FIGURES/6.

64. Muskovic, W., and Powell, J.E. (2021). DropletQC: improved identification of empty droplets and damaged cells in single-cell RNA-seq data. Genome Biol 22, 1–9. 10.1186/S13059-021-02547-0/FIGURES/2.

65. CellTypist | automated cell type annotation for scRNA-seq datasets https://www.celltypist.org/.

66. Elmentaite, R., Kumasaka, N., Roberts, K., Fleming, A., Dann, E., King, H.W., Kleshchevnikov, V., Dabrowska, M., Pritchard, S., Bolt, L., et al. (2021). Cells of the human intestinal tract mapped across space and time. Nature 2021 597:7875 597, 250–255. 10.1038/s41586-021-03852-1.

67. Van de Sande, B., Flerin, C., Davie, K., De Waegeneer, M., Hulselmans, G., Aibar, S., Seurinck, R., Saelens, W., Cannoodt, R., Rouchon, Q., et al. (2020). A scalable SCENIC workflow for single-cell gene regulatory network analysis. Nature Protocols 2020 15:7 15, 2247–2276. 10.1038/s41596-020-0336-2.

68. Yu, G., Wang, L.G., Han, Y., and He, Q.Y. (2012). clusterProfiler: an R Package for Comparing Biological Themes Among Gene Clusters. OMICS 16, 284. 10.1089/OMI.2011.0118.

69. Wu, T., Hu, E., Xu, S., Chen, M., Guo, P., Dai, Z., Feng, T., Zhou, L., Tang, W., Zhan, L., et al. (2021). clusterProfiler 4.0: A universal enrichment tool for interpreting omics data. The Innovation 2, 100141. 10.1016/J.XINN.2021.100141.

70. Liberzon, A., Subramanian, A., Pinchback, R., Thorvaldsdóttir, H., Tamayo, P., and Mesirov, J.P. (2011). Molecular signatures database (MSigDB) 3.0. Bioinformatics 27, 1739–1740. 10.1093/BIOINFORMATICS/BTR260.

71. Liberzon, A., Birger, C., Thorvaldsdóttir, H., Ghandi, M., Mesirov, J.P., and Tamayo, P. (2015). The Molecular Signatures Database (MSigDB) hallmark gene set collection. Cell Syst 1, 417. 10.1016/J.CELS.2015.12.004.

72. Nishimura, D. (2004). BioCarta. https://home.liebertpub.com/bsi 2, 117–120. 10.1089/152791601750294344.

73. Shao, X., Taha, I.N., Clauser, K.R., Gao, Y. (Tom), and Naba, A. (2020). MatrisomeDB: the ECM-protein knowledge database. Nucleic Acids Res 48, D1136–D1144. 10.1093/NAR/GKZ849.

74. Kaukonen, R., Jacquemet, G., Hamidi, H., and Ivaska, J. (2017). Cell-derived matrices for studying cell proliferation and directional migration in a complex 3D microenvironment. Nat Protoc 12, 2376–2390. 10.1038/NPROT.2017.107.

