## Supplemental item Table S1 for "Intercellular interaction between FAP fibroblasts and CD150 inflammatory monocytes mediates fibro-stenosis in Crohn’s disease"

### SUPPLEMENTAL ITEMS

**Table S1. Modified scoring system for histological assessment of inflammation in Crohn's disease, related to Figure S1B**

| Feature | Score |
|---|---|
| No increased inflammation | 0 |
| Inactive chronic inflammation (abnormal mucosal architecture, Pseudopyloric metaplasia and Paneth cell hyperplasia, mononuclear inflammatory cell infiltrate) | 1 |
| Mild active chronic inflammation (Neutrophils in lamina propria or epithelium without crypt abscesses) | 2 |
| Moderately active chronic inflammation (Crypt abscesses, destruction or loss of surface epithelium mixed with fibrin) | 3 |
| Erosions (Mucosal defect with muscularis mucosae still present) | 4 |
| Flat ulceration (Mucosal defect without residual muscularis mucosae) | 5 |
| Fissuring ulceration (Perpendicular into the deep submucosa or muscularis propria) | 6 |
| Fistula or abscess (Penetrating the bowel wall) | 7 |
