## Supplemental item Table S2 for "Intercellular interaction between FAP fibroblasts and CD150 inflammatory monocytes mediates fibro-stenosis in Crohn’s disease"

**Table S2. Fibrosis scoring system for histological assessment of fibrosis in Crohn's disease, related to Figure S1B**

| Layer | Feature | Score |
|---|---|---|
| Muscularis mucosa | Normal | 0 |
| | Thickening or splaying | 1 |
| | Downward extension (obliterative muscularization of the submucosa) | 2 |
| | Fusion with muscularis propria | 3 |
| Submucosa | Normal | 0 |
| | Thickening due to oedema | 1 |
| | Thickening due to fibrosis | 2 |
| | Upward extension of fibrosis to the mucosa layer | 3 |
| Muscularis propria | Normal | 0 |
| | Muscularis hyperplasia and thickening | 1 |
| | Collagen deposition (fibrosis) | 2 |
| | Fibrous septa through the layer | 3 |
| (Sub) serosa | Normal | 0 |
| | Collagen deposition | 1 |
| | Thickening | 2 |
