## Supplemental item Table S3 for "Intercellular interaction between FAP fibroblasts and CD150 inflammatory monocytes mediates fibro-stenosis in Crohn’s disease"

**Table S3. Clinical characteristics of patients included for scRNA-seq, related to Figure 1 and Figure S1**

| Characteristic | CD patients (n=10) | CRC patients (n=5) |
|---|---|---|
| General data | | |
| Age (y/o; mean ± SD) | 37.5 ± 11.2 | 67 ± 14.4 |
| Sex (M/F) | 3/7 | 4/1 |
| Disease duration (y; mean ± SD) | 10.1 ± 4.5 | N/D |
| Smokers (active/former/never) | 1/4/5 | N/D |
| Montreal classification | | |
| Age at diagnosis | | |
| A1 below 16 y/o | 0 | N/A |
| A2 between 17 and 40 y/o | 10 | N/A |
| A3 above 40 y/o | 0 | N/A |
| Location | | |
| L1 ileum | 8 | N/A |
| L2 colonic | 0 | N/A |
| L3 ileocolonic | 2 | N/A |
| L4 isolated upper disease | 0 | N/A |
| Behaviour | | |
| B1 non-stricturing, non-penetrating | 0 | N/A |
| B2 stricturing | 7 | N/A |
| B3 penetrating | 3 | N/A |
| P perianal disease modifier | 1 | N/A |
| Lab data | | |
| CRP (mg/L; mean ± SD) | 27.8 ± 27.1 | N/D |
| Medication naive | 1 | N/D |
| Medication at time of surgery | | |
| anti-TNFα | 2 | N/D |
| Ustekinumab | 4 | N/D |
| Vedolizumab | 1 | N/D |
| Steroids | 3 | N/D |

y/o, years old; y, years; N/A, not applicable; N/D, no data; CRP, c-reactive protein
